# OpenFISH enables integrated high-resolution spatial transcriptomics and metabolomics on a single tissue section

**DOI:** 10.1101/2025.08.19.671030

**Authors:** Xinyang Li, Yuan Huang, Shuo Wang, Ya Li, Fengting Jiang, Jiawei Gao, Yaran Yang, QingFeng Wu, Woo-ping Ge, Lihui Duan

## Abstract

Spatial transcriptomics enables *in situ* mapping of gene expression, yet no current platform provides single-cell, same-section integration with metabolomics, limiting direct links between transcriptional programsand metabolic phenotypesin native tissue. We present OpenFISH, a rapid, imaging-based spatial transcriptomics method operable on standard microscopes, requiring no proprietary hardware, and fully compatible with matrix-assisted laser desorption/ionization mass spectrometry imaging (MALDI–MSI). OpenFISH resolves hundreds of transcripts at subcellular resolution within 24 h and can be performed after MALDI–MSI, preserving metabolite distributions for cell-accurate co-registration on the same section. In mouse brain, integration with MALDI–MSI resolved metabolic heterogeneity at the level of individual cells. OpenFISH also quantified cell type–specific transcriptional activation of transposable elements after systemic lipopolysaccharide (LPS) challenge and detected disrupted spatial organization of D1 striatal neurons in Reeler mutants. Benchmarking showed performance comparable to or exceeding commercial platforms at ∼0.5% of per-sample cost. By enabling same-section, near-single-cell co-mapping of transcripts and metabolites in an accessible workflow, OpenFISH provides a scalable framework for high-content spatial multi-omics across neuroscience, immunology, cancer biology, and beyond.

## Introduction

Understanding how cellular identity and metabolic state are spatially organized is central to dissecting tissue function in health and disease^1–8^. Single-cell transcriptomics has transformed our view of cellular heterogeneity, but its dissociative nature erases spatial context. Spatial transcriptomics (ST) bridges this gap by enabling *in situ* mapping of gene expression within intact tissues^9–33^. However, current ST platforms—particularly those integrated with metabolomic imaging—remain limited by coarse spatial resolution, high cost, and complex instrumentation, restricting their use in spatial multi-omics^8^.

Matrix-assisted laser desorption/ionization mass spectrometry imaging (MALDI–MSI) provides label-free, spatially resolved maps of metabolites directly from tissue. Integrating MALDI–MSI with ST offers a powerful means to link transcriptional programs to metabolic phenotypes within their native microenvironment. Early integrations of Visium (50 µm) with MALDI–MSI (∼100 µm) demonstrate proof-of-principle for linking transcripts and metabolites, while leaving room to advance from multi-cell to single-cell resolution^8^. Other advances, including SpaceM, achieve single-cell MALDI metabolomics but are optimized for cultured cells and do not support simultaneous *in situ* transcriptomic profiling^7^.

We developed OpenFISH, a rapid, imaging-based spatial transcriptomics method that runs on standard microscopes, maps hundreds of transcripts within ∼24 h, and achieves cellular-resolution profiling. Crucially, OpenFISH is fully compatible with MALDI–MSI on the same tissue section and can be performed after MALDI–MSI. This same-section, near-single-cell integration directly links transcript-defined cell identities to local metabolite signals, enabling pathway-level and cell type–resolved analyses of metabolic heterogeneity that are not accessible to transcriptomics or metabolomics alone.

We demonstrate OpenFISH across diverse biological contexts. In mouse brain, integration with MALDI–MSI resolved metabolic heterogeneity at near-single-cell resolution. Following systemic lipopolysaccharide (LPS) challenge, OpenFISH quantified cell type–specific transcriptional activation of transposable elements. In the Reeler mouse, it detected disrupted spatial organization of D1 striatal neurons. Benchmarking against leading commercial platforms shows OpenFISH deliverscomparable or superiorperformanceat a fraction of the cost, establishing it as a broadly accessible and scalable framework for high-resolution spatial multi-omics.

## Results

### Design of OpenFISH

Spatial transcriptomics (ST) methods intended for broad laboratory use should be cost-effective, scalable, and operable on standard equipment, without requiring custom-built, high-resolution microscopy systems. Compatibility with other spatial omics modalities is also critical for enabling multi-layered molecular profiling *in situ*. For targeted ST approaches, flexibility in probe panel design and the ability to adapt cost-efficiently to different experimental contexts are key for large-scale adoption.

OpenFISH introduces a modular probe architecture that separates the padlock probe from gene-specific split probes, in contrast to existing rolling circle amplification (RCA)-based ST methods that rely on direct hybridization of costly, gene-specific padlock probes to target mRNA or cDNA **(Fig. 1a**). This design permits reuse of a pre-optimized padlock probes across different target panels and enables structural optimization to improve hybridization efficiency and tissue penetration **(Fig. 1a**). As in prior RCA-based implementations^34^, OpenFISH employs ReadOut (RO) probes incorporating only three of the four DNA bases to reduce secondary structure formation and enhance hybridization kinetics. Hybridization times are minimized to 1 h for padlock probes and 10 min for RO probes, shortening the workflow without loss of detection accuracy.

**Fig 1.**
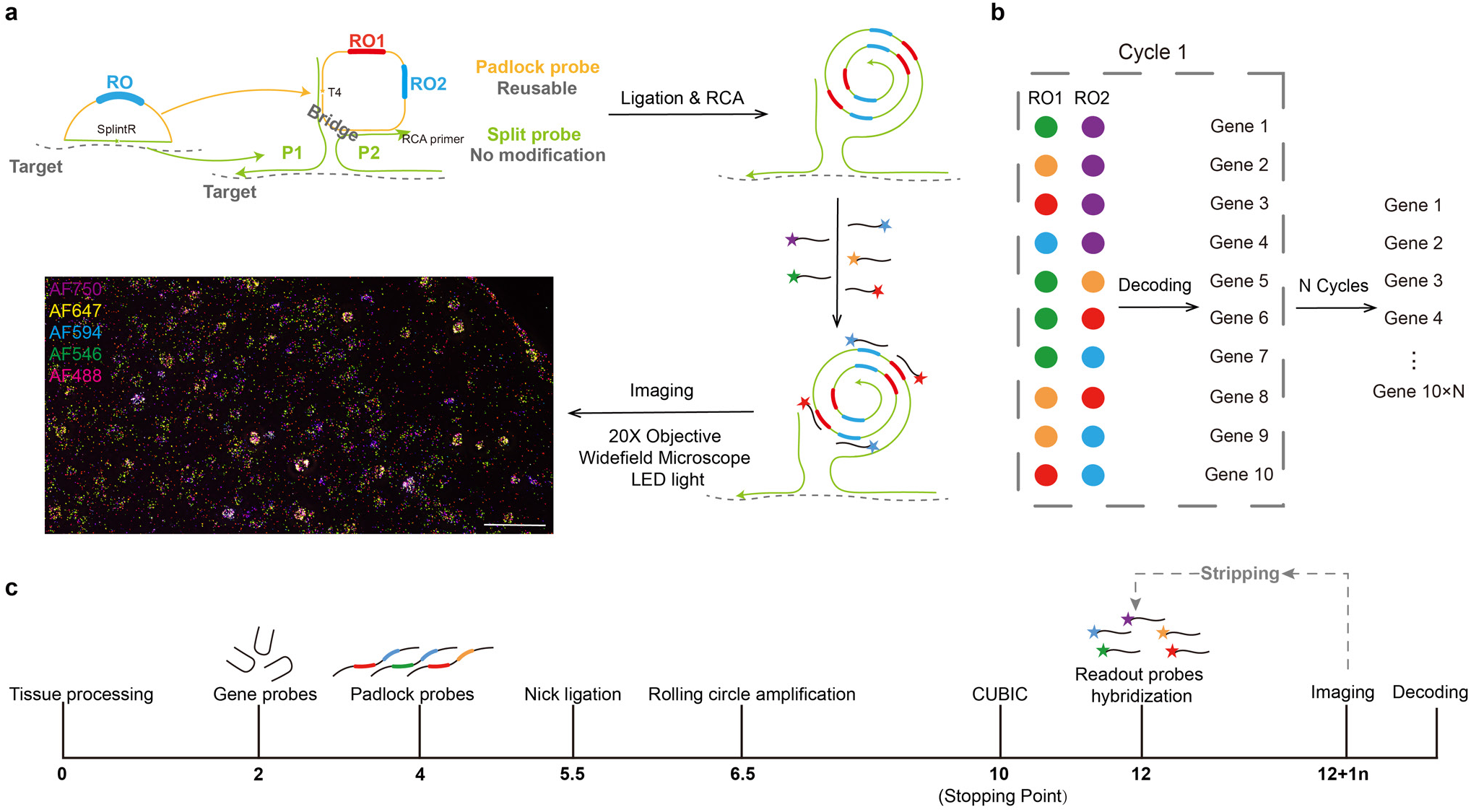
OpenFISH Design. **a,** Schematic of the OpenFISH process. Scale bar: 200 μm. **b,** OpenFISH encoding strategy. **c,** Timeline of OpenFISH bench work.

For high-throughput multiplexing, OpenFISH uses a simplified two-word encoding scheme derived from five fluorescent channels, excluding identical-word combinations to reduce background and improve decoding specificity (**Fig. 1b**). This approach avoids reliance on subpixel-level image registration, mitigating errors from sample drift and optical distortion during multi-cycle imaging and increasing computational efficiency. The wet-lab protocol requires ∼10 h, followed by ∼1 h per imaging round (**Fig. 1c**).

The pipeline was refined through iterative optimization of probe structure, target panel composition, noise-reduction coding, placeholder probe design to minimize signal crowding, CytoRNA staining to enhance cell segmentation, and incorporation of indium tin oxide (ITO) slide coating and tissue clearing—key steps enabling direct compatibility of OpenFISH with MALDI– MSI. Performance was validated by probe omission assays to confirm signal specificity, co-staining with corresponding target proteins to assess spatial concordance, and benchmarking against published datasets for quantitative evaluation (**Supplementary Note “Optimization of OpenFISH” and Supplementary Fig. 1–6**).

### Benchmarking of OpenFISH

We benchmarked OpenFISH against leading commercial spatial transcriptomics platforms to demonstrate comparable or superior sensitivity, spatial accuracy, and cell-type resolution, at a fraction of the cost and with unique compatibility for post-MALDI–MSI analysis. The mouse brain, with its extensively characterized anatomy and well-annotated gene expression at both single-cell and spatial scales, provides an ideal sample for benchmarking OpenFISH, allowing direct performance comparison with existing spatial transcriptomics datasets. A 110-gene panel was designed to capture a broad range of neuronal and non-neuronal cell types (**Fig. 2a**), and a false-positive control probe was included to quantify background signal. Computational modeling of the selected panel predicted a mean 96% classification accuracy (**Supplementary Fig. 7a**).

**Fig 2.**
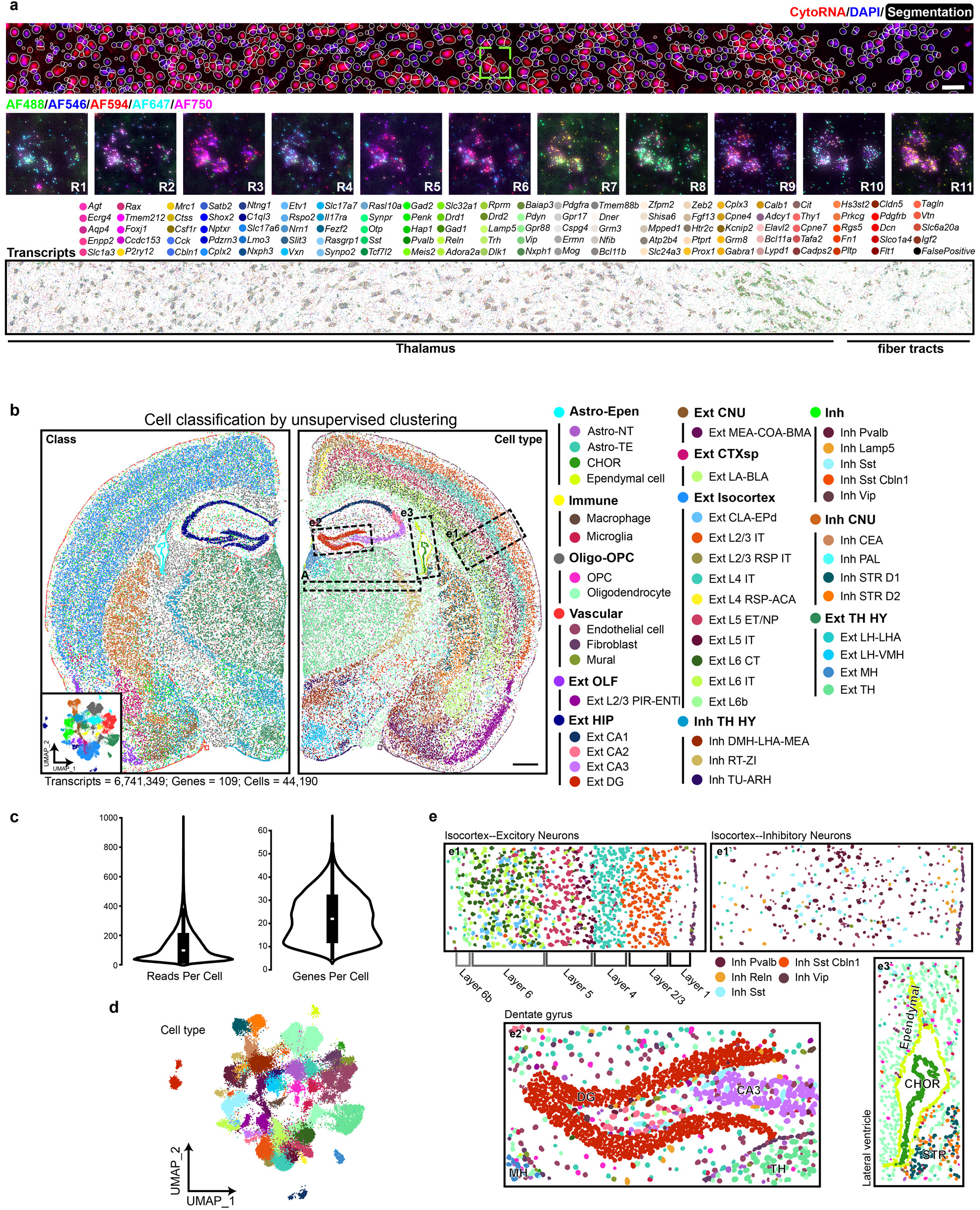
Performance of OpenFISH. **a,** Cellular segmentation achieved by the joint application of DAPI and CytoRNA (top) and raw signal image of highlighted ROI across 11 rounds (middle) and spatial distribution of decoded transcripts (bottom). Scale bar: 100 μm. **b,** The hemisected mouse brain was symmetrically presented as a whole brain. The segmentation results were plotted and annotated with class (left) and cell type (right). The half brain was classified into thirteen classes: Astro-Epen (Astrocytes and Ependymal cells); Immune; Oligo-OPC; Vascular; Ext OLF (Excitatory neurons, Olfactory areas); Ext HIP (Hippocampus); Ext CNU (Cerebral nuclei); Ext CTXsp (Cortical subplate); Ext Isocortex; Inh TH HY (Thalamus, Hypthalumus); Inh (Inhibitory neurons); Inh CNU; Ext TH HY. These classes can be further clustered into 44 distinct cell types. Astro-NT: Non-telencephalon Astrocytes; Astro-TE: Telencephalon Astrocytes; CHOR: Choroid plexus epithelial cells; OPC: Oligodendrocyte precursor cells; Mural: Pericytes and Smooth muscle cells; PIR: Piriform area; ENTl: Entorhinal area, lateral part; CA: Ammon’s horn; DG: Dentate Gyrus; MEA: Medial amygdalar nucleus; COA: Cortical amygdalar area; BMA: Basomedial amygdalar nucleus; LA: Lateral amygdalar nucleus; BLA: Basolateral amygdalar nucleus; CLA: Claustrum; EPd: Endopiriform nucleus; RSP: Retrosplenial area; IT: Intratelencephalic; ACA: Anterior cingulate area; CT: Corticothalamic; ET: Extratelencephalic; NP: Near-projecting; DMH: Dorsomedial nucleus of the hypothalamus; LHA: Lateral hypothalamic area; MEA: Medial amygdalar nucleus; RT: Reticular nucleus of the thalamus; ZI: Zona incerta; TU: Tuberal nucleus; ARH: Arcuate hypothalamic nucleus; CEA: Central amygdalar nucleus; PAL: Pallidum; LH: Lateral habenula; VMH: Ventromedial hypothalamic nucleus; MH: Medial habenula. Scale bar: 500 μm. **c,** Quality control of the OpenFISH data. **d,** UMAP visualization of cell types. **e,** Selected area shown in **b**, panel **e1** illustrates the distribution of different layers of excitatory neurons, while **e1’** displays the scattered inhibitory neurons from the same region as **e1**. Panel **e2** highlights the clear spatial distribution of dentate gyrus, whereas **e3** reveals the distinct spatial arrangement of astrocytes, ependymal cells and choroid plexus epithelial cells.

Applied to a coronal brain section, OpenFISH detected 6,741,349 transcripts across 44,190 cells (**Fig. 2b**), with a false-positive rate of 0.47%. The median transcript count was 97.5 per cell, with 22 median unique genes per cell (**Fig. 2c**). Forty-four distinct cell types were identified (**Fig. 2b, d, e**), and marker gene expression patterns were highly concordant with single-cell RNA sequencing (scRNA-seq) datasets (**Supplementary Fig. 8a**).

Detection efficiency—defined as the ratio of median transcripts per cell to panel size—was 0.894, exceeding that of Xenium (0.879) despite using fewer probes per target (**Supplementary Fig. 8b, c**). Clustering quality, quantified by Average Silhouette Width (ASW), was comparable to Xenium’s 248-gene panel (OpenFISH: 0.345, Xenium: 0.264, scRNA-seq: 0.412)(**Supplementary Fig. 8d**), demonstrating that OpenFISH resolves cellular heterogeneity with high precision using a more compact panel. Gene expression profiles were highly correlated across technical replicates (Pearson’s R = 0.97), confirming reproducibility (**Supplementary Fig. 8e**). To assess transcript-level accuracy independently of cell segmentation, we performed clustering directly on transcript coordinates. Without reference to scRNA-seq data, OpenFISH produced coherent clusters corresponding to major anatomical regions and cell types (**Supplementary Fig. 9a, b**).

Spatial accuracy was benchmarked against Visium HD, a high-resolution sequencing-based spatial transcriptomics platform. OpenFISH exhibited higher Moran’s I and lower Geary’s C values, indicating stronger spatial autocorrelation and improved preservation of native expression patterns (**Fig. 3a, b**). Mutual exclusivity correlation analysis showed transcriptional specificity comparable to Xenium and scRNA-seq, supporting accurate transcript assignment to distinct cellular compartments (**Fig. 3c–e**).

**Fig 3.**
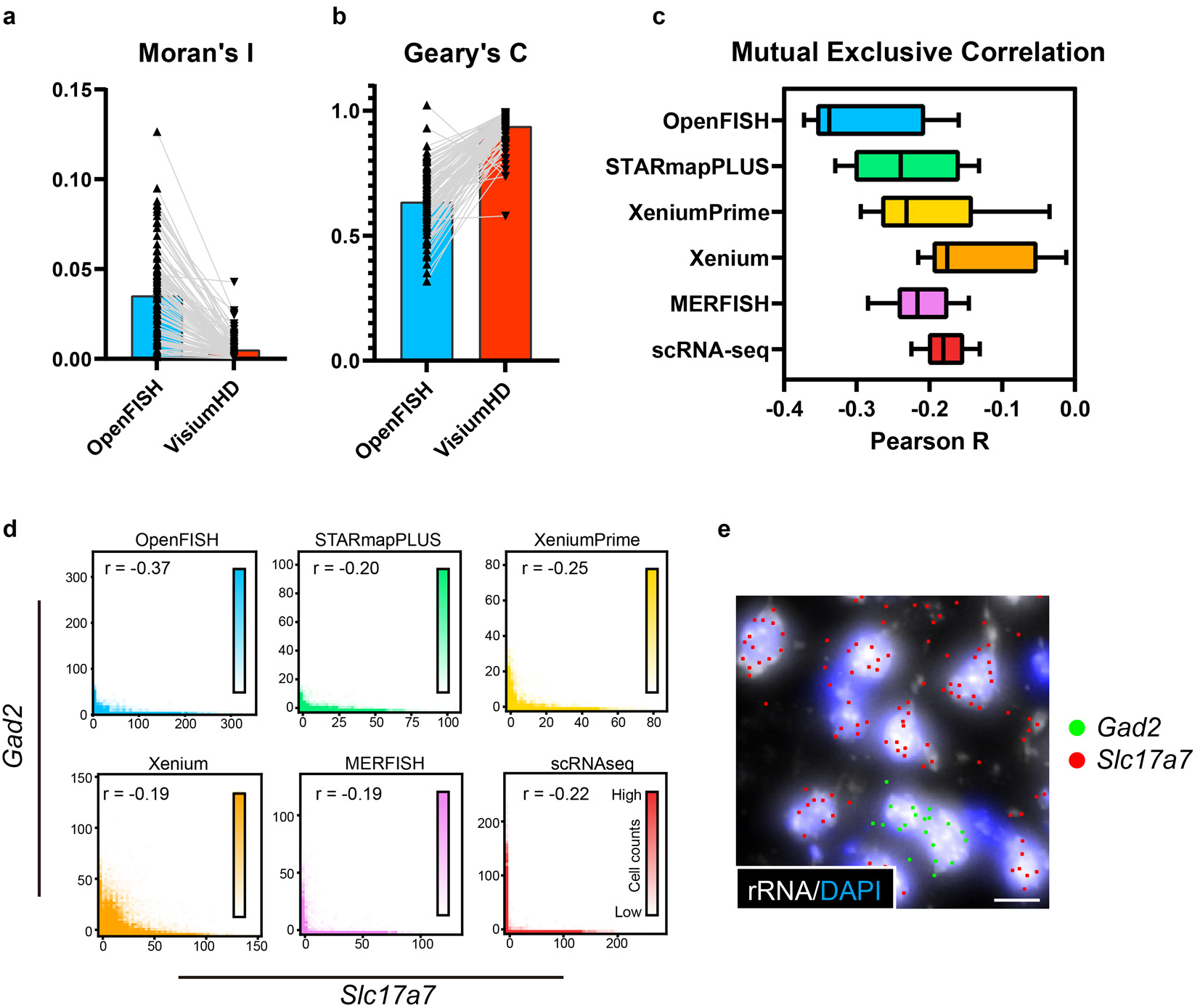
Benchmarking of OpenFISH Performance. **a,** Moran’s I for targeted genes. **b,** Geary’s C for targeted genes. **c,** Mutual exclusivity analysis. Selected genes are expected to show mutually exclusive expression in distinct cell types (Min to Max, n = 10, *Aqp4:Slc17a7*, *Aqp4:Gad2*, *Aqp4:Igf2*, *Aqp4:Slc17a6*, *Slc17a7:Gad2*, *Slc17a7:Igf2*, *Slc17a7:Slc17a6*, *Gad2:Igf2*, *Gad2:Slc17a6*, *Igf2:Slc17a6*). **d,** Expression plot of *Slc17a7* and *Gad2* across various methods per cell. **e,** OpenFISH example demonstration of *Slc17a7* and *Gad2* transcripts. Scale bar: 20 μm.

Together, these benchmarking results establish OpenFISH as a robust, high-fidelity spatial transcriptomics platform that matches or exceeds the performance of leading commercial systems while uniquely enabling integrated post-MALDI–MSI analysis in a cost-effective and accessible workflow.

### OpenFISH enables cell type–resolved mapping of metabolic, transcriptional, and structural alterations

#### Cell type–resolved metabolic landscapes via integration with MALDI–MSI

Systemic infection, modeled by LPS challenge, induces broad transcriptional and metabolic reprogramming across diverse brain cell types. While transcriptional responses have been extensively characterized^35^, the corresponding cell-type-specific metabolic adaptations remain less well defined.

We performed MALDI–MSI on the brains of saline- and LPS-treated mice at 20 µm spatial resolution—approaching the size of individual cells—and applied OpenFISH on the same tissue sections to map cell types across the brain (**Fig. 4a, b**). In total, we detected 1,290 distinct m/z species across the section, with an average of 918 species per pixel. And 80 median transcript count for 173,155 cells. Transcriptomic and metabolomic datasets were co-registered using an automated pipeline, with transcriptomic data aggregated to the same 20 µm resolution as MALDI–MSI, which improved the spatial precision of metabolite clustering (**Fig. 4c**).

**Fig 4.**
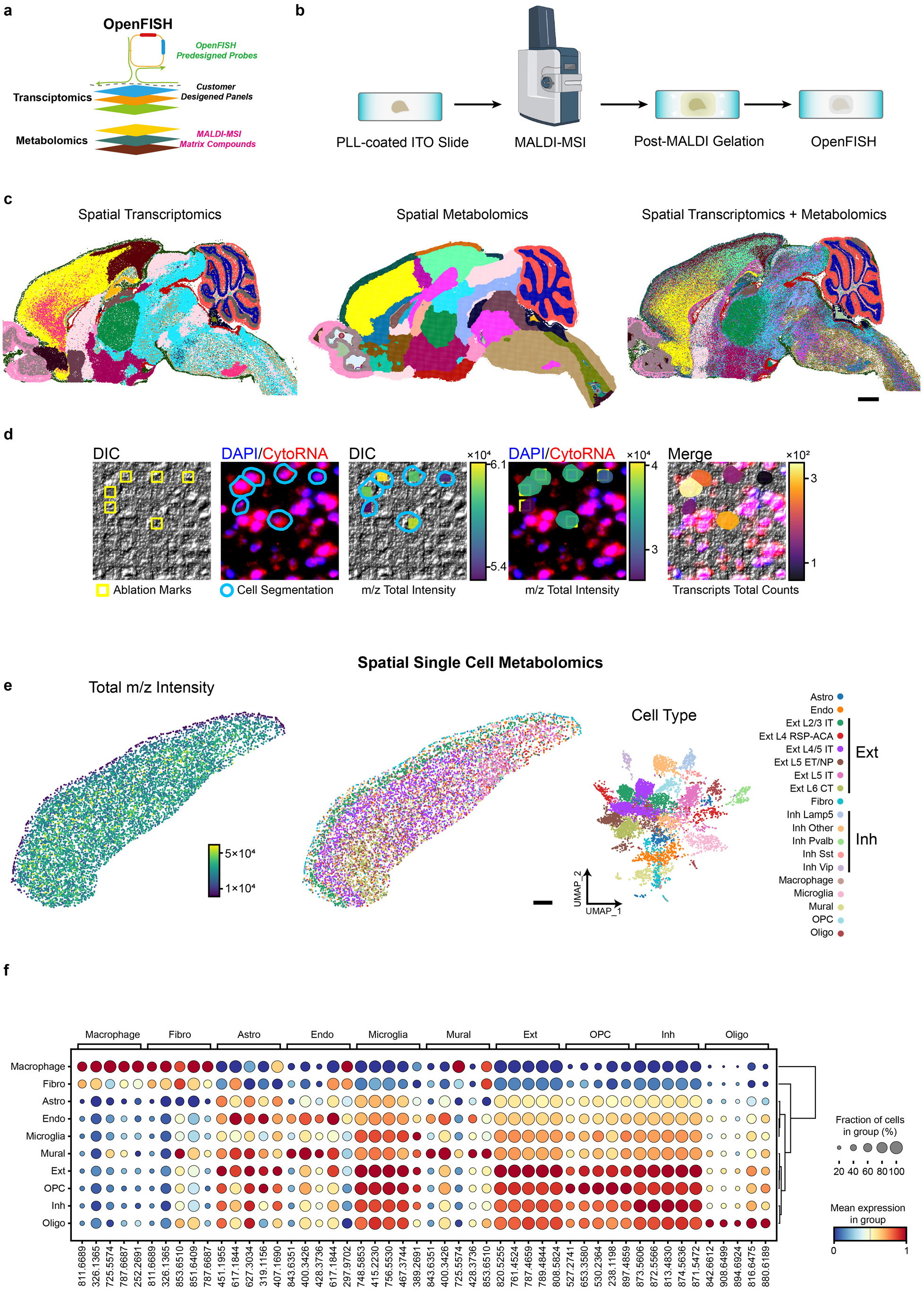
Integration of OpenFISH and MALDI-MSI in the same tissue. **a,** Schematic diagram of same tissue OpenFISH and MALDI-MSI. **b,** Experimental procedure of OpenFISH with MALDI–MSI. **c,** Clustering result using spatial transcriptomics modality only (left). Clustering result using spatial metabolomics modality only (middle). Clustering result using integrated spatial transcriptomics and metabolomics modalities (right). Scale bar: 1000 μm. **d,** Workflow for single-cell co-registration of spatial transcriptomics and metabolomics. **e,** Total m/z intensity distribution across cells in cortical region (left). Cell type annotation in cortical region (middle and right). Scale bar: 500 μm. **f,** Dotplot of differentially expressed metabolites m/z of each cell type in the cortical region.

To resolve metabolic landscapes at single-cell resolution, we focused on cortical regions where MALDI–MSI ablation marks were clearly identifiable in differential interference contrast (DIC) images, enabling accurate cell–pixel alignment (**Fig. 4d**). After filtering, 7,500 single cells were retained for analysis (**Fig. 4e**). Among all cell types, oligodendrocytes displayed highly distinctive metabolic profile, clearly separated from neurons, astrocytes, and other glial populations (**Fig. 4f**). The most strongly LPS-induced metabolite (m/z 650.4392; fold change = 1.68; **Supplementary Fig. 10a**) was putatively annotated as PoxnoPC, a truncated oxidized phosphatidylcholine within the OxPL class, which accumulates during inflammatory stress and can mediate both pro- and anti-inflammatory responses^36–38^. In cortex it widely distributed across cell types except fibroblasts, and reached highest abundance in excitatory neurons (**Supplementary Fig. 10a, b**). OpenFISH additionally identified LPS-reactive neurons using *Fos* probes, consistent with previously reported results^39–41^ (**Supplementary Fig. 10c**), and PoxnoPC was also enriched in these regions (**Supplementary Fig. 10d–f**).

### Cell type–specific transcriptional responses to inflammation

We next asked whether OpenFISH could resolve cell-type-specific transcriptional adaptations to inflammation. Recent evidenceindicatesthat transposable elements (TEs) can act as endogenous pathogen-associated molecular patterns (PAMPs) that trigger innate immune responses during inflammation^42^. We profiled brains fromthreepairs saline- and LPS-treated mice with a probe panel including probes targeting TEs (**Fig. 5a, b and Supplementary Fig. 11**). Median transcripts per cell and cell counts were: Saline1, 106.5 (58,536 cells); Saline2, 106.0 (51,998); Saline3, 106.5 (58,900); LPS1, 98.0 (52,619); LPS2, 111.5 (59,021); LPS3, 85.5 (55,184). In all LPS-treated animals, the ERV family member *MMVL30-int* was robustly upregulated in fibroblasts, showing a spatially widespread pattern with enrichment at the meninges and perivascular regions (**Fig. 5c, d**). Additional TE activation patterns showed clear cell-type specificity, including *RLTR4_MM-int* in endothelial cells and *RLTR6_Mm* in microglia. Differential expression analysis yielded concordant induction signatures across biological replicates (mixed-effects model, n=3 per condition): *MMVL30-int* in fibroblasts (β = 4.123, 95% CI 3.719–4.527, p < 0.001; between-sample variance = 0.044), *RLTR4_MM-int* in endothelial cells (β = 0.876, 95% CI 0.584–1.169, p < 0.001; 0.033) and *RLTR6_Mm* in microglia (β = 0.545, 95% CI 0.390–0.700, p < 0.001; 0.008). These estimates were consistent across animals, indicating high reproducibility of *in situ* quantification with OpenFISH.

**Fig 5.**
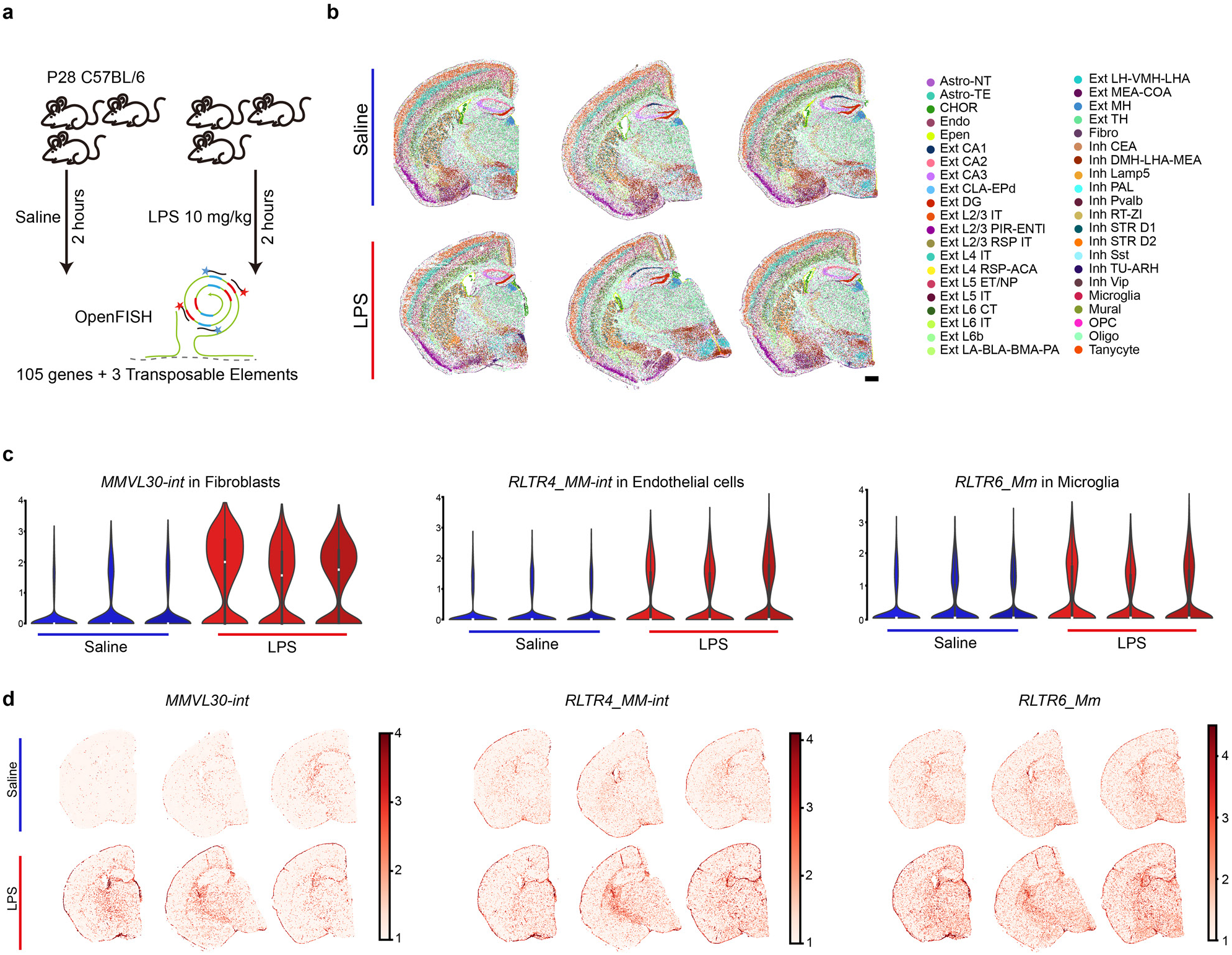
Cell type–specific transcriptional responses to inflammation. **a,** Schematic of the OpenFISH experimental design for detecting TE expression changes. **b,** Cell type composition and annotation in LPS-treated and saline-treated control samples. PA: Posterior amygdalar nucleus; PAL: Pallidum. Scale bar: 500 μm. **c,** Cell type–specific alterations in TE expression patterns following LPS treatment. **d,** Spatial alterations in TE expression patterns following LPS treatment.

### Detection of known and novel structural abnormalities in a neurodevelopmental model

Having established its ability to capture dynamic molecular responses, we evaluated OpenFISH for detecting fixed structural abnormalities. We applied OpenFISH to the Reeler mouse, a well-established model of *Reln* loss-of-function that disrupts neuronal migration^43–45^ and profiled two wild-type (WT) and two knockout (KO) brains. Median transcripts per cell and cell counts were: WT1, 74.0 (163,318 cells); WT2, 82.0 (136,407); KO1, 94.5 (138,279); KO2, 100.5 (113,881) (**Fig. 6a, b; Supplementary Fig. 12a, b**). OpenFISH faithfully recapitulated the previously reported disorganization of cortical (**Fig. 6c**) and hippocampal architecture (**Fig. 6d**), as well as cerebellar lamination (**Fig. 6e**), while also revealing additional, previously undescribed alterations in the striatum (**Fig. 6f, g**). In the cortex, excitatory subtypes from deep and superficial layers were thoroughly intermingled, with cell–cell neighbourhood analysis revealing rewired local networks (**Fig. 6c and Supplementary Fig. 12c, d**). The hippocampus exhibited pronounced dispersion of principal neurons, with clustering efficiency reduced by 12.3% in dentate gyrus and 15.9% in CA regions (**Fig. 6d and Supplementary Fig. 12e**). In the cerebellum, *Reln* knockout led to a marked reduction of excitatory neurons (Ext CB; *−*56%), accompanied by an increase in inhibitory neurons (Inh CB; +42%), astrocytes (+117%), and oligodendrocytes (+113%), as well as prominent Purkinje cell misplacement into inappropriate layers (**Fig. 6e and Supplementary Fig. 12f, g**). In the striatum, D1-type projection neurons were reduced by 36%, and the D1/D2 ratio decreased from 0.610 to 0.345 (−43%) (**Fig. 6g; Supplementary Fig. 12h, i**). To our knowledge, this has not been previously reported in Reeler. These data establish that OpenFISH can sensitively resolve both known and novel structural abnormalities at single-cell resolution, across multiple brain regions.

**Fig 6.**
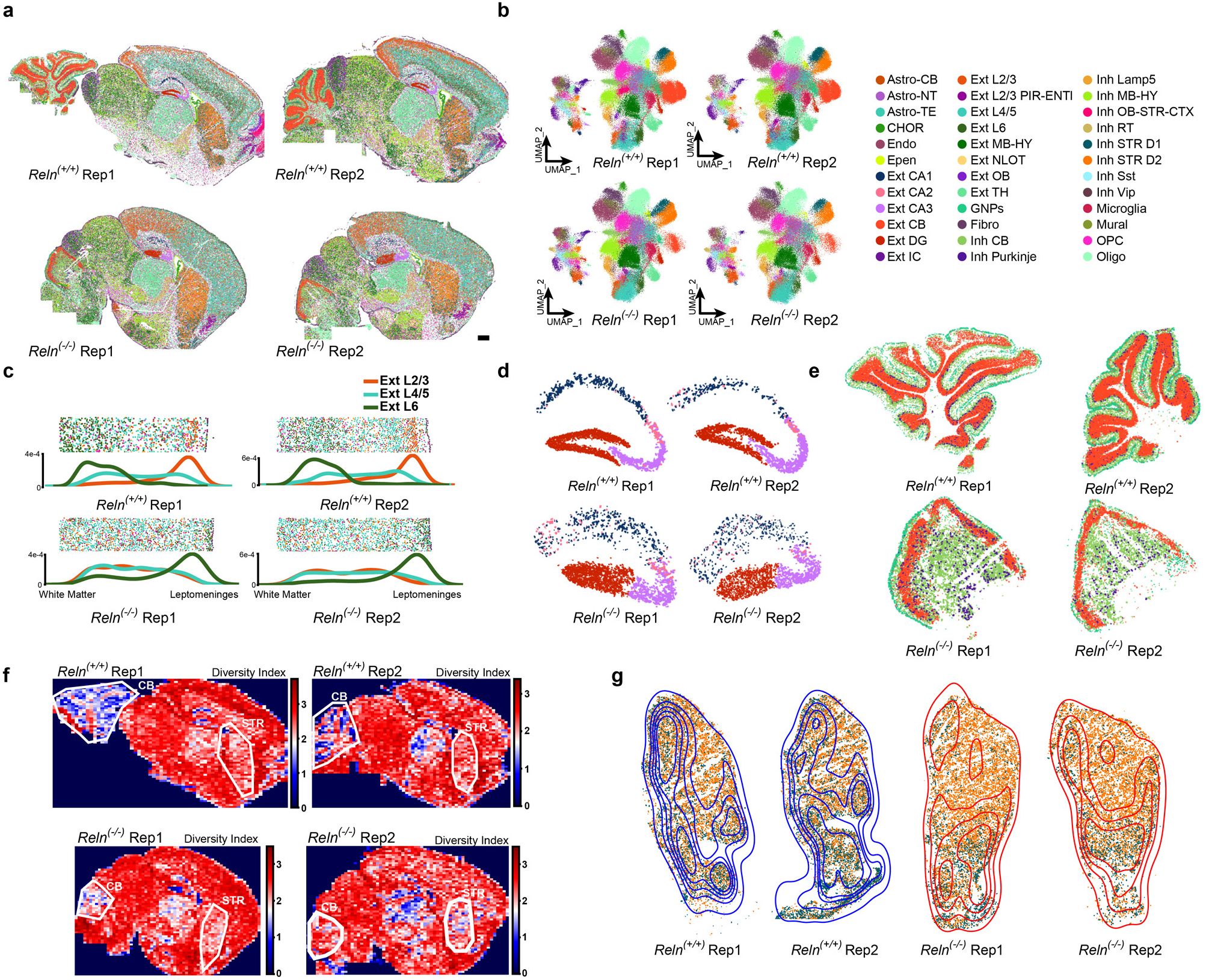
Structural abnormalities in *Reln*^(-/-)^ neurodevelopmental model. **a,** Spatial distribution of cell types in *Reln*^(+/+)^ and *Reln*^(-/-)^ samples, Scale bar: 1000 μm. **b,** UMAP projection with annotated cell types. IC: Inferior colliculus; CB: Cerebellum; MB: Midbrain; NLOT: Nucleus of the lateral olfactory tract; OB: Olfactory Bulb; GNPs: Granule Neuron Precursor. **c,** Cortical region: excitatory neuron density along the white matter-to-leptomeninges axis. **d,** Hippocampal region, highlighting Ext DG, Ext CA1, Ext CA2, and Ext CA3 subtypes. **e,** Cerebellar region across four samples. **f,** Global diversity index heatmap computed using MESA. **g,** Striatal region featuring Inh STR D1 and Inh STR D2 subtypes, with a Kernel Density Estimate (KDE) overlay based on Inh STR D1 spatial distribution.

Together, these results demonstrate that OpenFISH enables high-resolution mapping of cell type−resolved metabolic states, quantitative profiling of transcriptional responses, and sensitive detection of both known and novel structural abnormalities, providing a unified platform for dissecting architectural and functional changes across diverse disease models.

## Discussion

Spatial transcriptomics resolves gene expression within intact tissue, but integration with metabolomics—particularly matrix-assisted laser desorption/ionization mass spectrometry imaging (MALDI–MSI)—has been limited by spatial resolution, tissue handling, and cross-modality alignment, preventing single-section, near-single-cell multi-omics.

OpenFISH addresses these constraints by enabling imaging-based spatial transcriptomics after MALDI–MSI on the same tissue section. Optimization of ITO slide coating, hydrogel embedding, and tissue clearing rendered MALDI–MSI compatible with subsequent OpenFISH, and a streamlined co-registration pipeline aligned transcriptomic and metabolomic maps at near-single-cell resolution.

Applying this integrated workflow in a systemic LPS model allowed direct alignment of transcript-defined cell identities with local metabolite signals. The analysis revealed pronounced cell-type differences in metabolic profiles, with oligodendrocytes clearly separated from neurons, astrocytes, and other glia, while excitatory and inhibitory neurons exhibited largely similar signatures and identified LPS-associated specie (m/z 650.4392) enriched regions and cell types. These results show that same-section co-mapping can uncover biochemical patterns that are not discernible from transcriptomics or metabolomics alone.

Beyond multi-omics integration, OpenFISH provides a rapid, cost-effective spatial transcriptomics workflow operable on standard microscopes. Its modular probe architecture decouples gene-independent padlock probes from gene-specific split probes, reducing reagent cost and enabling rapid panel reconfiguration. The method typically requires ∼10 h of benchwork and ∼1 h per imaging round, facilitating inclusion of biological replicates. Benchmarking indicated performance comparable to or exceeding leading commercial platforms while operating at ∼0.5% (US$10) of per-sample cost (Table S2). OpenFISH reproduced known and identified additional structural features in the Reeler brain, quantified cell type–specific transcriptional activation of transposable elements in the LPS model, and enabled same-section integration with MALDI–MSI. While OpenFISH currently profiles 10 genes per imaging round, this targeted design maximizes detection fidelity and imaging efficiency for hypothesis driven studies. Ongoing improvements in encoding strategies, imaging throughput, and automated analysis are expected to expand its coding capacity and further enhance multi-omics integration.

In sum, OpenFISH establishes an accessible framework for high resolution, spatial multi-omics, enabling direct correlation of transcriptomic programs with metabolic phenotypes *in situ*. By combining technical accessibility, cost efficiency, and integrative power, it stands to democratize spatial multi-omics and accelerate discovery across neuroscience, immunology, cancer biology, and beyond.

## Methods

### Animals samples and ethics

All animal procedures used in this study were performed according to protocols approved by the Institutional Animal Care and Use Committee at the Institute of Genetics and Developmental Biology, Chinese Academy of Sciences. Mice were kept on C57BL/6 background and under a 12 h light–dark cycle with food and water provided ad libitum from the cage lid. They were under a temperature of 20– 25 °C and humidity of 30–70%. *Reln*-KO mice (Strain NO. T009126) were purchased from GemPharmatech (Nanjing, China).

### Panel Design

To design an optimal gene panel for delineating cell types in the mouse brain, we employed both automated and manual approaches. Initially, we tested various gene selection tools, using the Allen Brain Atlas scRNA-seq data as a reference^46^. We excluded certain cell subclasses that are only present in the olfactory bulb or cerebellum, with assistance from the Allen Brain Atlas MERSCOPE/MERFISH datasets^46^. Next, we eliminated lowly expressed genes (with an expression proportion within a cell type lower than 5%) and highly expressed genes (with total counts in the 99th percentile or above), which were then used as input for the panel design tools. We subsequently utilized the selected genes to re-cluster the scRNA-seq data, allowing us to compare the degrees of cell type separation in the newly projected embeddings. After conducting several benchmarks, we ultimately selected scGIST as our final tool. scGIST is a deep neural network-based tool that excels with small panel sizes^47^. Additionally, we reviewed published *in situ* hybridization and *in situ* sequencing data to select several well-known marker genes as the foundation for our panel. Specifically, we curated known excitatory neuron markers (*Slc17a7, Slc17a6*), inhibitory neuron markers (*Gad1, Gad2*), leptomeningeal cell markers (*Vtn, Slc6a20a*), and ependymal cell markers (*Tmem212, Foxj1*) from the literature. We prioritized these manually selected genes as input for scGIST. Finally, we ran scGIST for 200 epochs across five iterations, selecting the results that yielded the best accuracy and macro F1 score. In total, we selected 109 genes and one false-positive control for our final panel. To ensure an even distribution of genes across different cycles and channels, we employed a genetic algorithm to find the optimal gene combinations. Specifically, we first used MAGIC^48^ to impute the scRNA-seq reference expression data, which served as input for correlation calculations. We initialized the input with a population size of 300. For each gene combination, we sequentially assigned the genes to their respective cycles and calculated the Spearman coefficient within each cycle. The sum of all coefficients across all cycles was used as the fitness function. This process was iterated for a total of 500 generations. Similarly, within each cycle, we assigned genes to different channels using the same approach to avoid overcrowding, utilizing a population size of 127 and iterating for 300 generations. We then determined the number of probe pairs required for each gene. For most genes, two pairs of probes were sufficient, while a few required adjustments to the number of pairs for efficient detection. Following a pilot experiment, we may introduce placeholders for certain highly expressed genes or for genes that are unresolvable due to overcrowding. Detailed information of pair numbers and placeholders for each gene are listed in Table S3.

### Read Out (RO) Probes Design

To minimize potential secondary structures in rolling circle amplification (RCA) products and maximize hybridization efficiency, we first examined all 20 nt sequences composed solely of A, T, and C. The sequences were then filtered based on base composition (A: 10%–35%, T: 10%–35%, C: 40%– 60%), the presence of repeats (AAAA, TTTT, CCC), and the minimum free energy of secondary structures (using NUPACK^49^ default parameters, with a threshold of mfe > –0.5). The resulting RO probes were subjected to a BLASTn search against the mouse refseq_rna databases using the following parameters: blastn -task blastn-short -evalue 100 -strand minus. Probes aligning to the reference for more than 14 bp were excluded. The remaining candidates were handled similarly to the DeLOB^50^ procedure. Specifically, candidates were BLASTed against themselves with the parameters: blastn -task blastn-short -evalue 100 -strand both -max_target_seqs 10000. From this analysis, a graph was constructed in which each probe is represented as a node. An edge is drawn between two nodes if one probe aligns with another for more than 10 bp. A node was then randomly selected, and its edges and associated nodes were removed from the graph. This process was repeated with new random selections until no edges remained in the graph.

### Padlock Design

The padlock consists of one bridge sequence and two RO sequences, which were designed as described above. The bridge is a 30 nt sequence composed of A, T, C, and G. We first utilized seqwalk^51^ to generate 15 nt sequencing-level orthogonal sequences with the following parameters: seqwalk.design.max_orthogonality (500000, 15, alphabet = "ATCG", RCfree = True, GClims = (6, 14)). The generated sequences were then BLASTed against the refseq_rna database and the designed RO probes, with both thresholds set at 7 bp. Next, the half bridges were combined to form complete 30 nt bridges. The DeLOB method was employed to ensure the orthogonality of the bridges, following the same criteria as before. The results were subsequently BLASTed against refseq_rna for final specificity evaluation. In each round, five RO probes were randomly selected from the available RO probes. Since each padlock requires two different RO probes, a total of 10 padlocks were designed for each round. For each RO combination within a cycle, a bridge was randomly chosen from the previously designed bridge database. The complete padlock was then assembled, and the secondary minimum free energy (mfe) was assessed using NUPACK with the parameters: material = "dna", celsius = 37, sodium = 0.075, magnesium = 0.01, with a threshold of mfe set at –1. The entire padlock, along with gene probe 1, gene probe 2, and the target mRNA, was used as input for a complex tube check in NUPACK with same parameters as above. We aimed for an optimal padlock configuration where the ends of the complex are exposed.

### Gene Probe Design

For a given gene, if its sequence is not provided, the corresponding mRNA will be automatically downloaded from NCBI. Each gene probe consists of a 30 nt region that targets the mRNA, a 2 nt spacer (AT), and a 14 nt region that binds to the bridges on the padlock. Gene probe 1 and gene probe 2 are designed with a two-nucleotide spacing between their target regions. The probe generation process involves four main steps: (1) Basic Filter; (2) Specificity Filter; (3) Binding Efficiency Filter; and (4) Evaluation. A user-friendly notebook is available to facilitate the probe design. Initially, candidate targets are filtered based on their base complexity (Shannon entropy < 1). Probes containing repeats (such as GGGGG or CCCCC) or with a melting temperature (Tm) lower than the expected 47 °C are excluded. Tm is calculated using Biopython packages with the following parameters: nn_table=MeltingTemp.R_DNA_NN1, dnac1=20, selfcomp=True, Na=390, and Mg=0.0. Tm is then corrected using the chem_correction function with parameters: fmd=30 and fmdmethod=2. Next, the probes are subjected to a BLASTn search with the parameters: blastn -task blastn-short -evalue 10 - strand minus. Any nonspecific alignments with a binding part Tm greater than 37 °C are excluded. The complete probe will then be assembled using the provided padlock sequence. If a blacklist file is provided, the probes will be BLASTed against this blacklist to ensure specificity. Binding efficiency is calculated using NUPACK, defined as the probability of forming the target complex in a test tube compared to the probability of forming both the target and background complexes. The background complex consists of mRNA and the secondary structure probabilities of probe 1 and probe 2. Finally, all candidates are evaluated based on GC percentage, Tm, non-target Tm, the difference in Tm of probe 1 and probe 2, and binding efficiency. A score is assigned to each probe pair after evaluation. The final probes are synthesized by Sangon using HAP purification.

### CytoRNA Design

18S rRNA sequence was downloaded from NCBI and sent into Primer3 for primer design. Primer expected length is set to 30 nt to decrease the possibility of unspecific binding to any other artificial sequences.

### TE (Transposable Element) Probe Design

Raw sequencing data were obtained from the Gene Expression Omnibus (GEO) under accession number GSE112436. Read alignment was performed using Cell Ranger (v7.0.1) with default parameters against the mm39 reference genome. The resulting aligned BAM files were processed using SoloTE^52^ for transposable element (TE) quantification, with TE annotations derived from the UCSC RepeatMasker (rmsk) BED file. Gene expression matrices (incorporating both gene and TE subfamily features) from LPS-treated and saline-treated samples were harmonized using Harmony. Following manual cell type annotation, we employed COSG^53^ to identify differentially expressed TE subfamilies across cell types. To characterize TE expression at locus-level resolution, we examined expression patterns of individual loci within selected TE subfamilies. The most highly and variably expressed TE loci, along with their corresponding genomic sequences, were selected for probe design as follows. Besides normal probe design procedure, potential off-target effects were assessed through Bowtie2 alignment to evaluate probe specificity.

### Lipopolysaccharide (LPS) Treatment

4-week-old mice were intraperitoneally injected with a single dose of LPS (Escherichia coli, serotype O111:B4, Sigma-Aldrich, Cat# L2630-25MG; 10 mg/kg). Control animals received an equal volume of saline. Mice were sacrificed 2 h post-injection.

### Sample preparation

The mice were euthanized, and fresh brain tissues were rapidly excised. The brains were embedded in OCT, and the tissues were rapidly frozen in isopentane pre-chilled with dry ice, followed by storage at −80 °C. For MALDI–MSI and OpenFISH on same slides, brain tissues were embedded in pre-cooled FSC22 or without embedding. Then the tissues were frozen on dry ice followed by storage at −80 °C. For tissue sectioning, mouse brains were transferred to a cryostat and sectioned into 10 µm thick slices, with the chamber temperature set to −20 °C and the specimen head temperature set to −10 °C. The slices were attached to microscope slides or PLL-treated ITO slides and stored at −80 °C until use. 1 mL 0.1 mg/mL poly-L-lysine (Biotopped Cat# Top0668) in H2O was applied onto the ITO slides for 1 h and thoroughly washed using DEPC H2O three times. Coated slides were then air dried for further use.

### OpenFISH protocol

The section was fixed with 4% (w/v) PFA (Electron Microscopy Sciences, Cat# 15710) at room temperature (RT) for 20 min, and then washed twice with PBS for 1 min each. The tissue was permeabilized in 0.5% Triton X-100 (Merck millipore, Cat# 648463) for 15 min at RT and washed two times with PBS for 1 min each before hybridization. The sample was incubated with blocking buffer (2× SSC, 1% Tween 20, 0.02% BSA, 0.5 μg/μL Salmon Sperm DNA sheared (Invitrogen, Cat# AM9680), 0.2 μg/μL yeast tRNA (Invitrogen, Cat# AM7119) and 0.4 U/μL Murine RNase inhibitor (Vazyme, Cat# R301-03)) at RT for 10 min and then washed two times with 0.5× SSCT buffer (0.5× SSC, 0.05% Tween 20 and 0.4 U/μL Murine RNase inhibitor) for 1 min each at 42 °C. Assemble a 2- mm-thick nanogrid tape with a scissor-cut hole and transparent film to form a sealed chamber that isolates the tissue from the outside environment.

The sample was incubated with gene hybridization buffer (2× SSC, 1% Tween 20, 0.02% BSA, 30% formamide (Sigma-Aldrich, Cat# F9037), 0.2 μg/μL yeast tRNA, 0.4 U/μL Murine RNase inhibitor and gene probes at 20 nM per oligo) at 42 °C for 90 min and then washed two times with 30% washing buffer (2× SSC, 30% formamide and 0.4 U/μL Murine RNase inhibitor) for 15 min each at 42 °C. Next, the sample was incubated with padlock hybridization buffer (2× SSC, 1% Tween 20, 0.02% BSA, 0.2 μg/μL yeast tRNA, 0.4 U/μL Murine RNase inhibitor and padlock probes at 20 nM per oligo) at 37 °C for 60 min and then washed two times with 2× SSC buffer (2× SSC and 0.4 U/μL Murine RNase inhibitor) for 5 min each at 37 °C. Finally, the sample was incubated with T4 ligase buffer (1× T4 DNA ligase buffer, 0.02% BSA, 7 U/μL T4 DNA Ligase (TaKaRa, Cat# 2011B) and 0.4 U/μL Murine RNase inhibitor) at RT for 60 min. After ligation, the sample was washed twice with 20% washing buffer (2× SSC, 20% formamide and 0.4 U/μL Murine RNase inhibitor) for 10 min at 37 °C and then washed three times with PBST (0.05% Tween 20 in PBS) for 1 min each at RT. The sample was incubated with RCA buffer (1× Phi29 buffer, 250 µM dNTPs, 0.5 U/μL Phi29 polymerase (Vazyme, Cat# N106-02) and 0.02% BSA) at 30 °C for 3.5 hours and then washed twice with PBST for 1 min each at RT. Finally, CUBIC- HL (TCI, Cat# T3781) was applied to the sample for 60 min at RT. Following this treatment, the sample was washed three times with PBS for 1 min at RT. Next, the sample was treated with CUBIC-R (TCI, Cat# T3741) for 10 min at RT to restore the refractive index, after which it was washed three more times with PBS for 1 min each. The slide was now ready for RO hybridization and imaging.

### MALDI**–**MSI

Slides were vacuum dried for 30 min. 2,5-Dihydroxybenzoic acid (DHB, Sigma-Aldrich, 15 mg/mL dissolved in 90% acetonitrile) was sprayed onto the tissue using TM-sprayer (60 °C, 20 passes, solvent flow rate of 100 μL/min, pressure of 9 psi and drying time between passes of 5 s). Tissue sections were imaged at 20 μm lateral resolution using timsTOF fleX MS system. Laser power was set to 90% and data were collected over the m/z 50–1300. Immediately after MALDI–MSI, slices were stored at −80 °C until use.

### Post-MALDI OpenFISH

The slides were taken out from −80 °C and equilibrate to RT. The slides were immersed in prechilled 70% ethanol for 30 s, followed by immersion in prechilled 100% ethanol for 30 s on ice. Subsequently, the slices were fixed with 4% PFA in PBS at RT for 20 min, and then washed twice with PBS for 1 min each. The tissue was permeabilized with 0.5% Triton X-100 and 1 μg/mL DAPI at RT for 15 min and washed two times with PBS for 1 min each. The tissue was immersed in Fluoromount-G (SouthernBiotech, Cat# 0100-01) and covered with clean coverslips for DIC and DAPI imaging. Afterwards, the coverslip was gently removed and tissue was washed with stripping buffer (80% formamide, 2× SSC and 0.1% Tween 20 in H_2_O) for 5min at 37 °C. After two washes using PBS, the slices were blocked at RT for 10 min in blocking buffer (2× SSC, 1% Tween 20, 0.2 μg/μL yeast tRNA, 0.02% BSA, 0.4 U/μL Murine RNase inhibitor and 2 μM polyA anchor probe). The polyA anchor probe had a mixture of DNA and locked nucleic acid (LNA) nucleotides (/5Acryd/TTTTT+TTTTT+TTTTT+TTTTT+TTTTT+TTTTT) where T+ is a locked nucleic acid and /5Acryd/ is a 5’-acrydite modification. After blocking, the slices were washed three times with PBST and incubated in 20 mM MOPS at RT for 30 min, followed by incubation in 20 mM MOPS buffer containing 1 mg/mL MelphaX^54^ and 0.1 mg/mL Acryloyl-X, SE (Invitrogen, Cat# A20770) at 37 °C for 1 h. The slices were then washed three times with PBST for 5 min each. To cast the tissue-hydrogel hybrid, the slices were first incubated with monomer buffer (2× SSC, 4% (v/v) 19:1 acrylamide:bis-acrylamide (Macklin, Cat# A853856), 0.2% (v/v) tetramethylethylenediamine [TEMED] (Mreda, Cat# M1651)) at room temperature for 30 min. Following incubation, the polymerization mixture (2× SSC, 4% (v/v) 19:1 acrylamide:bis-acrylamide, 0.2% (v/v) TEMED, and 0.2% (w/v) APS) was added to the center of the slices and covered with Gel Slick-coated coverslips. The polymerization process was conducted at RT under a nitrogen atmosphere for 90 min, followed by three washes with PBST for 5 min each. Afterwards, the slices were digested at 37 °C for 10 min in digestion buffer (2× SSC, 0.5% Triton X-100, 2% (v/v) sodium dodecyl sulfate [SDS] (Sigma-Aldrich, Cat# L3771), 1 mg/mL proteinase K (Thermo Scientific, Cat# EO0491), 0.8 U/μL Murine RNase inhibitor), and then washed three times with PBST for 5 min each. Subsequently, the OpenFISH protocol was performed as previously described.

### OpenFISH Imaging

Each cycle began with treating the sample with stripping buffer at 37 °C for 3 min two times, followed by 20% washing buffer (2× SSC, 20% formamide in H_2_O) washing for 1 min. The samples were incubated with RO probe hybridization buffer (2× SSC, 20% formamide, 0.02% BSA, 2 μg/mL DAPI, readout probes at 2 nM per oligo and readout blocker probes at 20 nM per oligo) at RT for 15 min. The samples were washed by 20% washing buffer for 3 min at RT and then immersed in Fluoromount-G for imaging. For hydrogel-embedded samples, each cycle began with treating the sample with stripping buffer at 37 °C for 6 min three times. And 20% washing buffer supplemented with 10 mM ascorbic acid was used after RO hybridization.

Images were acquired using Zeiss Imager.Z2 microscopy with X-Cite XYLIS LED, and a 20× objective (NA 0.8). We utilized five fluorescence channels: AlexaFluor 488, AlexaFluor 546, AlexaFluor 594, AlexaFluor 647, AlexaFluor 750 and a DAPI channel. Eleven cycles of imaging were performed to detect 109 genes and an extra cycle of imaging was performed to record CytoRNA. An initial ROI was defined in Zen during the first imaging round. For subsequent rounds, the ROI was repositioned after a quick DAPI scan to minimize tile-to-tile displacement.

### OpenFISH with concurrent immunostaining

OpenFISH was performed as described above. Subsequently, the sample was washed three times for 5 min each with PBS at RT, and incubated in blocking solution (5% BSA, 0.5% Triton X-100 in PBS) at RT for 60 min. The sample was then incubated with primary antibody (Rabbit anti-Somatostatin, Phoenix Pharmaceuticals, 1:1000) in blocking solution at 4 °C overnight, washed three times for 5 min each with PBS at RT, and incubated with secondary antibody (AlexaFluor 488-conjugated AffiniPure Goat anti-Rabbit-IgG (H+L), Jackson Lab, 1:1000) and DAPI (2 μg/mL) in blocking solution at RT for 1 h. The sample was washed three times for 5 min each in PBS at RT and mounted in Fluoromount-G for imaging.

### OpenFISH Raw Image Processing

Raw image files in .czi format were first combined into a maximum projection along the z-axis using the Z-stack projection function in Zen. Next, the images were deblurred with the following parameters: strength = 0.5, radius = 3, and sharpness = 0. After this, tiles were exported from each channel in each cycle, with the CytoRNA cycle exported without deblurring. The subsequent processing was performed in a compact Jupyter notebook using Python. Tiles were clipped to remove overexposed areas, and BaSiC^55^ was employed to eliminate camera shadows.

### OpenFISH Stitching and Registration

Using the DAPI channel as a reference, each cycle was registered to the first cycle using Euler transform in SimpleITK. The transformation matrix obtained from this step was then applied to correct the alignment of other channels, also addressing any drift caused by the microscopy hardware. Following this, the reference DAPI image from the first cycle was stitched into a complete image using m2stitch^56^, and global coordinates were assigned to the other cycles. After stitching, the entire image underwent further processing using adaptive histogram equalization to reduce intensity attenuation near the edges of the tiles.

### OpenFISH Segmentation

The preprocessed DAPI channel was segmented using StarDist^57^ with the following parameters: prob_thresh = 0.5, nms_thresh = 0.4, trained_model = "2D_versatile_fluo", and sigma = 2.5. DAPI segmentation was then expanded for 5 μm. The preprocessed CytoRNA channel was segmented using the Cellpose3 denoise model with parameters: model_type = "cyto3", restore_type = "deblur_cyto3", diameter = 45, flow_threshold = 1, and cellprob_threshold = -6. The segmentation results from Cellpose3^58^ were then geometrized using code from Sopa with minor modification for parallel processing. Finally, the segmentation results for DAPI and CytoRNA were resolved for conflicts using custom code. Specifically, DAPI segmentation masks were initially expanded by a 5-μm buffer. CytoRNA segmentation polygons were given priority over DAPI polygons in cases of overlap. For each intersection between a CytoRNA polygon and a DAPI polygon, the following criteria were applied: (1) If the overlapping area exceeded 30% of the DAPI polygon area, the DAPI polygon was excluded. (2) If the overlapping area exceeded 70% of the CytoRNA polygon area, the CytoRNA polygon was discarded as it likely represented background signal. (3) In all other cases, the intersecting boundary was used to split the polygons, generating two new distinct regions.

### OpenFISH Decoding

Stitched and registered images for each channel in each cycle were processed using RS-FISH^59^ via a Fiji macro. The following parameters were applied: anisotropyCoefficient = 1.00, ransac = RANSAC, sigmaDoG = 1.20, supportRadius = 3, inlierRatio = 0.1, maxError = 0.75, min_number_of_inliers = 6, initial = 8, final = 20, bsMethod = "RANSAC on Median", bsMaxError = 0.05, and bsInlierRatio = 0.1. Different channels had slightly varying thresholdDoG values: 0.0055 for AlexaFluor 488, 0.0060 for AlexaFluor 546, 0.0065 for AlexaFluor 594, 0.0055 for AlexaFluor 647, and 0.0060 for AlexaFluor 750. For some situations, Spotiflow^60^ was applied for spots detection using pre-trained model "hybiss" with parameters prob_thresh = 0.40, min_distance = 1, exclude_border = True, scale = None, subpix = True, peak_mode = "fast", normalizer = "auto". Spotiflow outputs were further filtered by intensity. Then the spots’ probabilities were sent as "intensity" for decoding. Subsequently, the RS-FISH or Spotiflow results were processed with PoSTCode^61^, with minor modifications. Specifically, we constructed a K-D Tree for spots within each channel. In a single cycle, the nearest spots between two channels were identified, and their corresponding intensities were adjusted using a Gaussian distribution with a mean of 0 and a standard deviation of 1. Subsequently, a matrix was constructed as input for PoSTCode, and after execution, multi-use spots were deduplicated using the Union-Find algorithm. The final decoded spots were filtered to exclude those with a nearest neighbor distance of less than 30 pixels. Each cell was then aggregated for spots that fell within the polygon of the cell segmentation. Additionally, the related raw images, segmentation results, decoded spots, and aggregated AnnData were integrated into a standard SpatialData^62^ object for downstream analysis. We also exported an Explorer file for convenient data inspection.

### OpenFISH Data Analysis

Brain slides were annotated using hierarchical clustering. Initially, raw data were filtered by cell segmentation area to remove abnormal cells, retaining those with an area, in pixels, larger than 600 or within three times the mean area. Counts for each gene were then adjusted according to the placeholder percentage. Next, cells with a false-positive rate of less than 1% were kept, followed by further filtering with the criteria: min_genes = 5, max_counts = 1000, and min_counts = 20. For Transposable elements related samples, criteria: min_genes = 2, max_counts = 900, and min_counts = 10 were used. For *Reln* related and MALDI samples, criteria: min_genes = 2, max_counts = 1000, and min_counts = 10 were used. We then normalized data by cell area and library size. After log1p processing, we employed the Leiden algorithm to cluster the cells, and differentially expressed genes for each cluster were calculated using COSG. For MALDI samples, CellLENS^63^ was used for spatially-aware clustering. Each cluster was manually inspected based on the spatial distribution of cells and gene expression. For clusters that appeared to contain multiple subtypes, we performed additional Leiden clustering on those specific clusters. Cell type results of each slide were imported into SpatialData and used for plotting.

### Segmentation-free spot clustering

The OpenFISH output was modified to serve as the input for FICTURE^64^, with the following parameters used: ficture run_together --all --mu-scale 3.077 --major-axis X --n-factor 30. Resulting coordinates were transformed to the original global coordinate system by: (1) dividing by 100, (2) adding the OFFSET value (specified in the third line of the result file), and (3) multiplying by 3.077. The scaled coordinates and their associated topic assignments were subsequently imported into SpatialData for visualization and further spatial analysis.

### Spatial autocorrelation and mutual exclusive analysis

Moran’s I was calculated using the moranfast R package, while Geary’s C was computed using the squidpy.gr.spatial_autocorr function in Squidpy^65^. Ten pairs of common genes were selected for mutual exclusivity analysis: (*Aqp4*, *Slc17a7*), (*Aqp4*, *Gad2*), (*Aqp4*, *Igf2*), (*Aqp4*, *Slc17a6*), (*Slc17a7*, *Gad2*), (*Slc17a7*, *Igf2*), (*Slc17a7*, *Slc17a6*), (*Gad2*, *Igf2*), (*Gad2*, *Slc17a6*), and (*Igf2*, *Slc17a6*). Pearson residual was calculated between each gene pairs for each method as the indicator for mutual exclusive evaluation.

### Cluster benchmark

The Average Silhouette Width (ASW) was calculated using the scib.me.silhouette function from the scib^66^ package on the UMAP embedding. The label key was set to either cell type or equivalent classification for each method. Nearest mutual information (NMI) was calculated using the scib.metrics.nmi function from scib package on the UMAP embedding.

### MALDI Data Processing

Raw m/z data were normalized using root mean square (RMS) normalization in SCiLS Lab software to stabilize the data and remove batch effects. For all spots within the same experimental samples, spectral peaks were aligned^67^, and the processed data were exported in imzML format. The datasets were then imported into Python using the pyimzML package, and an expression matrix was constructed using the spatialMETA package. To enable spatial registration, raw spot coordinates were first scaled to approximate their real-world dimensions (0.325 μm for one pixel), and a mean intensity image was generated for alignment. Both the DIC (differential interference contrast) image and the mean intensity image were loaded into napari for manual registration. The resulting affine transformation matrix was applied to the scaled spot coordinates to align them with the reference image. Aligned coordinates were used to generate a grid of square polygons (13 μm length, 20 μm spacing). These polygons served as spatial bins to aggregate spot-level data, analogous to cell segmentation, allowing for coarse assessment of two-modality integration performance. For finer registration, square polygons were exported as Fiji ROI (Region of Interest) files, and each polygon’s position was manually adjusted based on ablation traces to ensure precise alignment. The final registered MALDI data were used to derive single-cell-level spatial metabolomics. For m/z intensity assignment to single cells, our approach closely followed the SpaceM^68^ method with minor modifications. The sampling area for each cell was defined as the intersection between a cell segmentation mask and a square polygon. Sampling specificity was calculated as the ratio of the sampling area to the total intersection area of the square polygon, while the cell area was defined by the segmentation mask. Only cells meeting stringent criteria (sampling area/cell area > 0.45 and sampling specificity > 0.90) were retained for downstream analysis. Finally, for a given metabolite, intensities were adjusted based on the relative sampling area:

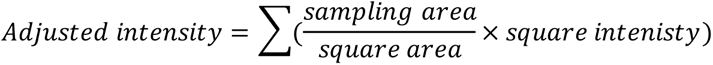

### MALDI**–**OpenFISH Integration Analysis

Transcripts data were assigned to each square polygon as previously mentioned. The square-level OpenFISH data were filtered with the criteria: min_genes = 3, max_counts = 250, and min_counts = 10. The MALDI ion data were first cleaned by removing background ion signals (Table S4). Those background signals were partially manual selected according to ion image, and other ion m/z fell below 200 or exceeds 1000 were excluded. Then we filtered squares using the criteria: min_ions = 400, max_intensity= 60000, and min_intensity = 8000. 110 highly variable ions were selected for MALDI data. After, each cell in OpenFISH data and MALDI data were normalized to have a total counts of 110 and concatenated into a single anndata object. CellLENS was applied to get an initial result of clustering. The clustering results were sent to MENDER^69^ for another round of spatially-aware clustering with parameters: nn_mode= "radius", scale = 6, radius = 60. OpenFISH data and MALDI data were also separately processed with this procedure.

### Reeler mouse Data Analysis

Samples were first preprocessed and annotated as previously mentioned. For each ROI, included cells were sent as input for MESA^70^ to comprehensively assess the structural alteration. Scale = 64 was applied to all ROIs. Other parts follow the tutorial of MESA with annotated cell type as feature to calculate cell type frequency and cell-cell co-occurrence frequency. For Hippocampus region, clustering efficiency was calculated using networkx package. Briefly, a network was constructed with a 100-pixel threshold. And clustering coefficient was calculated using networkx.average_clustering function.

## Code availability

All codes for recreating the analyses in the figures have been deposited at GitHub at https://github.com/Lixinyoung/openfish-maldi.

## Data availability

All the sequences used in this work are listed in Table S1.

Source data provided with this paper will be deposited in Zenodo.

Public data used in this manuscript are available via the following links.

Allen Brain Atlas scRNAseq data and MERSCOPE/MERFISH data were downloaded from: https://portal.brain-map.org/atlases-and-data/bkp/abc-atlas;

Xenium mouse brain data was downloaded from: https://www.10xgenomics.com/datasets/fresh-frozen-mouse-brain-for-xenium-explorer-demo-1-standard;

XeniumPrime5k mouse brain data was downloaded from: https://www.10xgenomics.com/datasets/xenium-prime-fresh-frozen-mouse-brain; STARmapPLUS data was downloaded from: https://singlecell.broadinstitute.org/single_cell/study/SCP1375/integrative-in-situ-mapping-of-single-cell-transcriptional-states-and-tissue-histopathology-in-an-alzheimer-disease-model;

## Acknowledgements

We gratefully acknowledge Dr. Jason G. Cyster and Dr. Xiang Yu for comments on the manuscript. We thank members of the Duan laboratory for comments and suggestions. The workwas supported by National Key Research and Development Program of China (2024YFA1803400), and National Natural Science Foundation of China (32371023).

## Author contributions

L.D. conceived the project. L.D., X.L., Y.H., S.W., Y.L. designed experiments. X.L. designed and implemented the data analysis code. X.L., Y.H., S.W., Y.L. performed experiments and analyzed data. F. J., andf J.G. performed the MALDI-MSI analysis. Y. Y. performed the other experiments. Q. W. and W-P. G.provided assistance in experimental design and data analysis. L.D., X. L. and Y. H. wrote the manuscript. All authors reviewed and edited the manuscript.

## Competing interests

The authors declare that they have applied for a patent related to this work.

## Correspondence and requests for materials

should be addressed to Lihui Duan.

