## Supplemental File 1 for "OpenFISH enables integrated high-resolution spatial transcriptomics and metabolomics on a single tissue section"

### Supplementary Note: Optimization of OpenFISH

OpenFISH is an open, low-cost spatial transcriptomics platform that integrates modular probe design, streamlined encoding, and optimized segmentation to deliver single-cell-resolution mapping directly on MALDI–MSI-processed tissue sections, overcoming the resolution, cost, and accessibility limitations of existing commercial systems.

#### Molecular design and validation of OpenFISH

In OpenFISH, unmodified split probes (P1 and P2) anneal to target mRNA at their 3′ and 5′ terminus, respectively. The padlock probe hybridizes to the 5′ terminus of P1 and the 3′ terminus of P2, generating a nick site within the DNA-DNA hybrid. Ligation is catalyzed by T4 DNA ligase (**Fig. 1a**), which offers higher specificity than SplintR ligase. Rolling circle amplification (RCA) initiates at the 3′ terminus of P2, ensuring that P1, P2, and the padlock probe are all necessary for amplification (**Supplementary Fig. 1a**). Ligation efficiency peaked when the nick site was located at the middle position within the 14 bp of the P1 5′ terminus. (**Supplementary Fig. 1b**). Specificity was confirmed through two approaches: (1) co-staining OpenFISH-detected *Sst* transcripts with the corresponding protein antibody (**Supplementary Fig. 1c**), and (2) comparing cortical layer marker distributions with publicly available spatial transcriptomic atlas, which showed high concordance in spatial localization (**Supplementary Fig. 1d**).

#### Probe length optimization for low-abundance targets

Conventional RNA detection methods such as HCR typically employ 20-nt target-binding split probes, but detection of low-abundance transcripts is often inefficient. For example, *Gjb2* expression in the meninges was detectable only when no fewer than 10 probe pairs were used. Extending the target-binding region to 30 nt and using only four probe pairs, markedly increased per-cell signal density and produced results comparable to the commercial Xenium platform, which used eight probes for the same gene (**Supplementary Fig. 2a, b**). These findings indicate that OpenFISH maintains high detection efficiency with fewer probes, reducing costs and simplifying probe panel design.

#### Encoding strategy and noise suppression

To maximize multiplexing capacity while minimizing secondary structure, each padlock probe carries two readout binding sites, enabling simultaneous detection of up to 15 genes per imaging round with five fluorophores (**Fig. 1b**). During validation, single-color spots were detected despite the two-color encoding expected for true signals. These monochrome spots were identified as noise, likely arising from misclassification of background fluorescence or imaging artifacts by high-sensitivity spot detection algorithms. To improve specificity, all single-color signals were excluded from decoding and analysis, retaining only dual-color-encoded signals. This refinement enabled reliable detection of 10 targets per imaging round (**Fig. 1b**).

#### Imaging efficiency

Although OpenFISH detects 10 targets per imaging round, its robust amplification and sparse signal distribution enable imaging with a standard 20× objective on an epifluorescence microscope, eliminating the need for high-resolution optics. This configuration supports rapid acquisition of large tissue areas; for example, imaging a half-coronal mouse brain section (0.6 cm × 0.6 cm × 10 μm) requires only ~30 minutes.

**Probe and panel design optimization**

Hybridization efficiency in spatial transcriptomics can be constrained by probe secondary structure, which slows hybridization kinetics and reduces tissue penetration. To address this limitation, OpenFISH employs target-independent padlock and readout probes designed for minimal secondary structure. Orthogonal artificial sequences were first generated using SeqWalk and DeLOB, then filtered with NUPACK to remove candidates with low hybridization kinetics or high secondary structure probability (**Supplementary Fig. 3a, b**).

For panel design, single-cell RNA sequencing (scRNA-seq) data was integrated with non-targeted spatial transcriptomics datasets. Genes showing spatial correlation *in situ* but not in scRNA-seq were prioritized, as they represent different spatially organized cell types. The gene list was refined using scGIST to maximize classification accuracy and macro F1 score (**Supplementary Fig. 3c, Supplementary Fig. 7a**). Expression level and spatial distribution were then used to allocate genes across imaging rounds via a genetic algorithm to minimize spatial overlap and signal saturation.

**Signal detection and error correction**

Spot detection and decoding were performed using RS-FISH and PoSTcode, with fluorescence channel intensity serving as the primary decoding criterion (**Supplementary Fig. 3d**). OpenFISH employs a dual-color encoding scheme where each transcript is labeled with two distinct readout probes. Additionally, the physical distance between paired spots was incorporated into PoSTcode, with spot intensity adjusted using a Gaussian attenuation function, allowing recovery of drifting spots and filtering of mismatches. To reduce overcrowding by high-abundance transcripts, a “placeholder padlock” lacking 5′ phosphorylation was introduced; it hybridizes to split probes but remains unligated, thereby limiting signal overcrowding. The final transcript counts were adjusted according to the padlock-to-placeholder padlock ratio(**Supplementary Fig. 4**). A false-positive control probe targeting a non-existing sequence was included to monitor background.

**Enhanced cell segmentation with CytoRNA staining**

Accurate cell segmentation in dense or morphologically complex tissues is hindered by the limited cytoplasmic information provided by DAPI staining. Expanding DAPI masks or assigning spots to the nearest nucleus may introduce segmentation errors and exclude cells lacking visible nuclei but containing cytoplasmic RNA. Although polyA staining can capture cytoplasmic boundaries, its low signal intensity limits practical use. We therefore developed CytoRNA staining, which simultaneously labels ribosomal RNA (rRNA) and messenger RNA (mRNA) by using rRNA-binding probes with flanked polyA sequences, followed by Oligo-dT-Cy3 staining. CytoRNA produced brighter and more continuous cytoplasmic signals than polyA staining, improving the ability to depict cell boundaries (**Supplementary Fig. 5a**).

Segmentation was performed with Cellpose3 for CytoRNA and StarDist for DAPI signals, followed by integration of CytoRNA boundaries with expanded DAPI masks. This approach increased the number of high-confidence cells, improved transcript retention per cell, and reduced contamination (**Supplementary Fig. 5b, c**).

#### Data processing and output format

Following spot detection and cell segmentation, transcripts were assigned to individual cells, and extracellular transcripts were excluded. Transcripts of the same gene with nearest-neighbor distances exceeding a predefined threshold were removed. The final datasets were stored as SpatialData objects, including raw images, segmentation polygons, decoded spots, and aggregated AnnData matrices, enabling integration with standard spatial transcriptomics analysis pipelines.

#### Integration of OpenFISH with MALDI–MSI on the same tissue section

Matrix-assisted laser desorption/ionization mass spectrometry imaging (MALDI–MSI) enables label-free, *in situ* biomolecule quantification in fresh-frozen tissue sections. Integrating spatial transcriptomics with MALDI–MSI allows direct correlation of gene expression with spatially resolved metabolic states at single-cell resolution. However, direct fluorescence imaging after MALDI–MSI was hindered by strong autofluorescence, resulting in poor signal-to-noise ratios (SNR) and precluding high-resolution transcript detection (**Supplementary Fig. 6a**).

To overcome this limitation, we developed a post-MSI hydrogel-embedding and tissue-clearing protocol to enhance optical clarity and SNR. Following MSI acquisition, matrix residues were removed with ethanol, tissues were fixed with paraformaldehyde and permeabilized with Triton X-100, and nucleic acids were anchored to the hydrogel matrix using MelphaX and polyT anchors. Subsequent protein digestion and lipid removal improved probe accessibility and reduced background fluorescence.

A major technical challenge was bubble formation and hydrogel delamination from ITO slides (**Supplementary Fig. 6b**), which frequently caused tissue deformation and registration errors. Pre-coating ITO slides with poly-L-lysine (PLL) markedly reduced bubble formation and prevented delamination without compromising MALDI–MSI performance (**Supplementary Fig. 6c**), thereby enabling consistent downstream imaging and accurate data alignment.

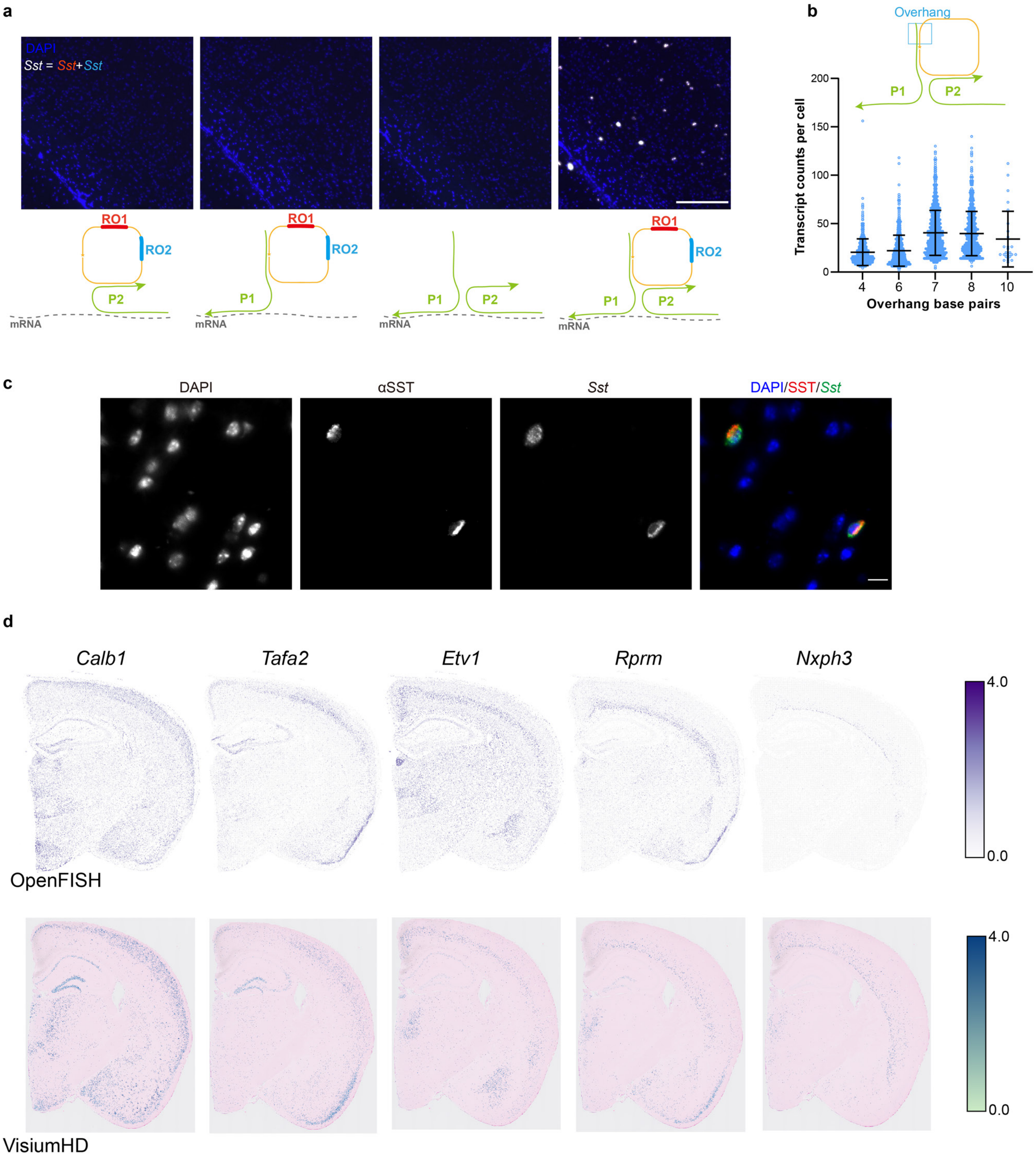

a
DAPI
Sst = Sst+Sst
RO1
RO2
P1
P2
mRNA
b
Overhang
Transcript counts per cell
Overhang base pairs
c
DAPI
αSST
Sst
DAPI/SST/Sst
d
Calb1
Tafa2
Etv1
Rprm
Nxph3
OpenFISH
VisiumHD
4.0
0.0

**Supplementary Fig. 1: Validation of OpenFISH**. **a,** The four fluorescence images showed *Sst* (red + blue = grey) mRNA and cell nuclei (blue) labeling in mouse brain sections, illustrating the following hybridization conditions from left to right: (1) P2 and padlock probe, (2) P1 and padlock probe, (3) P1 and P2, and (4) P1, P2, and padlock probe. Scale bar: 200 μm. **b,** The number of detected transcripts (*Sst*) per cell was quantified under conditions where the 3' end of the padlock probe and the 5' end of P1 exhibited varying degrees of complementarity, with base pair matches of 4, 6, 7, 8, and 10 nt. **c.** Left to right: DAPI (nuclei), immunofluorescence staining of SST protein (red), OpenFISH detection of *Sst* mRNA (green), merged channels. Scale bar: 10 μm. **d,** (top) Spatial plots of decoded features expression from a validation OpenFISH experiment. (bottom) The RNA patterns of individual genes extracted from Visium HD.

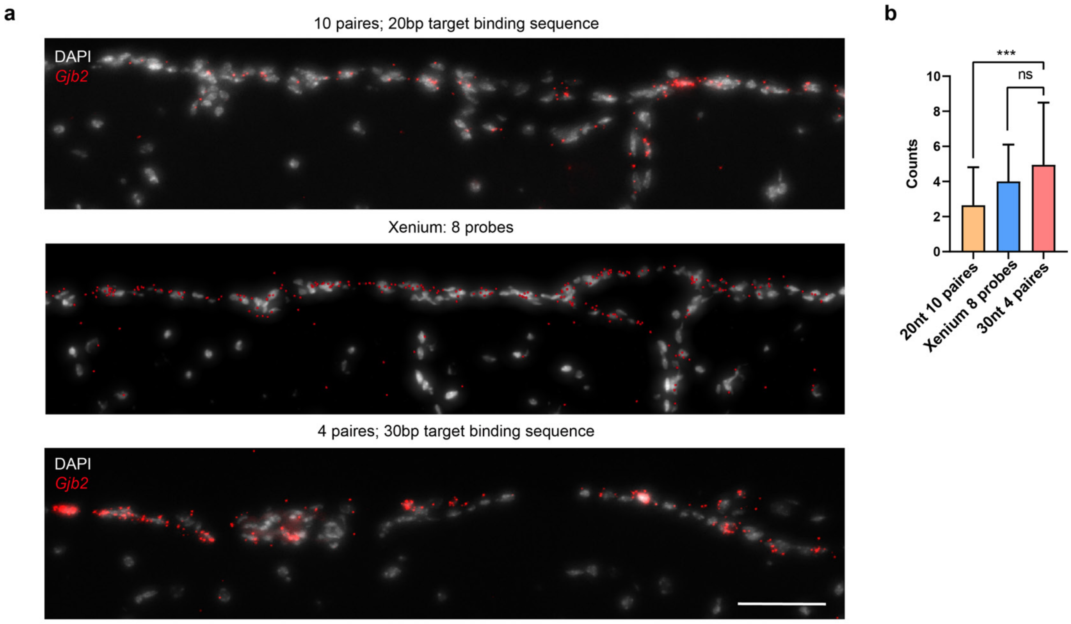
a
10 paires; 20bp target binding sequence
DAPI
Gjb2
Xenium: 8 probes
4 paires; 30bp target binding sequence
DAPI
Gjb2
b
***
ns
Counts
0
2
4
6
8
10
20nt 10 paires
Xenium 8 probes
30nt 4 paires

**Supplementary Fig. 2: Probe Length Optimization of OpenFISH**. **a,** The raw OpenFISH images and the processed Xenium images show *Gjb2* (red) mRNA and nuclei (grey) labeling within the mouse brain tissue sections. Optimization of gene probes was conducted using 10 pairs of gene probes (each probe binding to a 20 nt segment of mRNA) and 4 pairs of gene probes (each binding to a 30 nt segment). Xenium used probes targeting 8 distinct sites on the *Gjb2* mRNA. Scale bar: 50 μm. **b,** The quantification of *Gjb2* transcripts detected within each cell (left panel) (***p < 0.001, two-tailed unpaired Student's *t*-test, mean with SD).

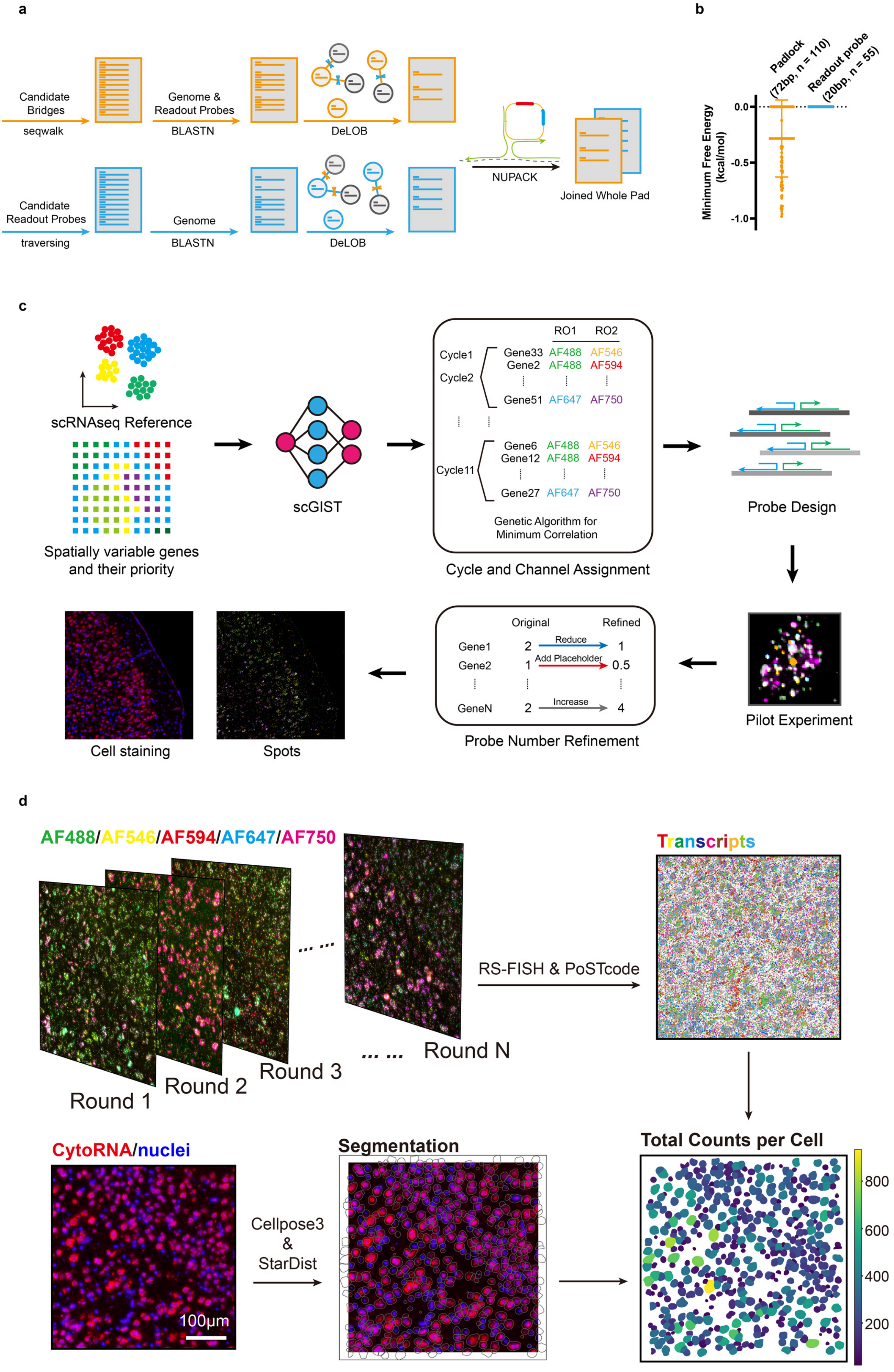

a
Candidate Bridges
seqwalk
Genome & Readout Probes
BLASTN
DeLOB
Candidate Readout Probes
traversing
Genome
BLASTN
DeLOB
NUPACK
Joined Whole Pad
b
Padlock (72bp, n = 110)
Readout probe (20bp, n = 55)
Minimum Free Energy (kcal/mol)
0.0
-0.5
-1.0
c
scRNAseq Reference
Spatially variable genes and their priority
scGIST
RO1 RO2
Cycle1
Cycle2
Gene33 AF488 AF546
Gene2 AF488 AF594
Gene51 AF647 AF750
Cycle11
Gene6 AF488 AF546
Gene12 AF488 AF594
Gene27 AF647 AF750
Genetic Algorithm for Minimum Correlation
Cycle and Channel Assignment
Probe Design
Pilot Experiment
Original Refined
Gene1 2 Reduce 1
Gene2 1 Add Placeholder 0.5
GeneN 2 Increase 4
Probe Number Refinement
Cell staining
Spots
d
AF488/AF546/AF594/AF647/AF750
Round 1
Round 2
Round 3
Round N
RS-FISH & PoSTcode
Transcripts
Total Counts per Cell
800
600
400
200
CytoRNA/nuclei
100μm
Cellpose3 & StarDist
Segmentation

**Supplementary Fig. 3: Computation Pipeline of OpenFISH. a,** Diagram of artificial sequence design. Candidate bridges were generated using SeqWalk, and the alignment tool BLASTn was employed to exclude sequences that exhibited cross-reactivity with the mouse genome. The remaining bridges were processed through the DeLOB algorithm to obtain the final orthogonal bridges. Readout probes were generated in a similar manner, with the exception of the candidate generation step, since RO sequences comprise only three bases, allowing all possible combinations to be enumerated. The final bridges and ROs were combined to create the complete padlock sequence. NUPACK was utilized for secondary structure inspection. **b,** The minimum free energy statistical diagrams of padlock (n = 110) and RO (n = 55) (Mean with SD). **c,** Flow chart of panel design. Candidate spatially variable genes were selected from published studies and assigned a priority score. A curated scRNA-seq reference, along with the prioritized genes, was input into scGIST. Multiple iterations of scGIST were conducted, and the best results were selected. A genetic algorithm was employed to assign genes to different cycles and channels. The assigned genes were then used for gene probe design. In addition to prior knowledge, a pilot experiment was conducted to determine the final number of probe pairs for each gene. **d,** Decoding pipeline of OpenFISH. A custom Jupyter Notebook pipeline was developed for efficient data processing. Raw tiles from each cycle were preprocessed to eliminate image abnormalities. After registration and stitching, RS-FISH was employed for spot detection. Detected spots were reconstructed and sent to PoSTcode for maximum-likelihood-based decoding. CytoRNA cycle images were also registered and stitched to the global coordinate system. To enhance cell recovery and computational efficiency, StarDist was utilized to segment only the DAPI channel, while Cellpose3 was used to segment the CytoRNA channel. The final results were merged and conflicts resolved, resulting in transcripts being aggregated into cells. A SpatialData object was generated for convenient downstream analysis.

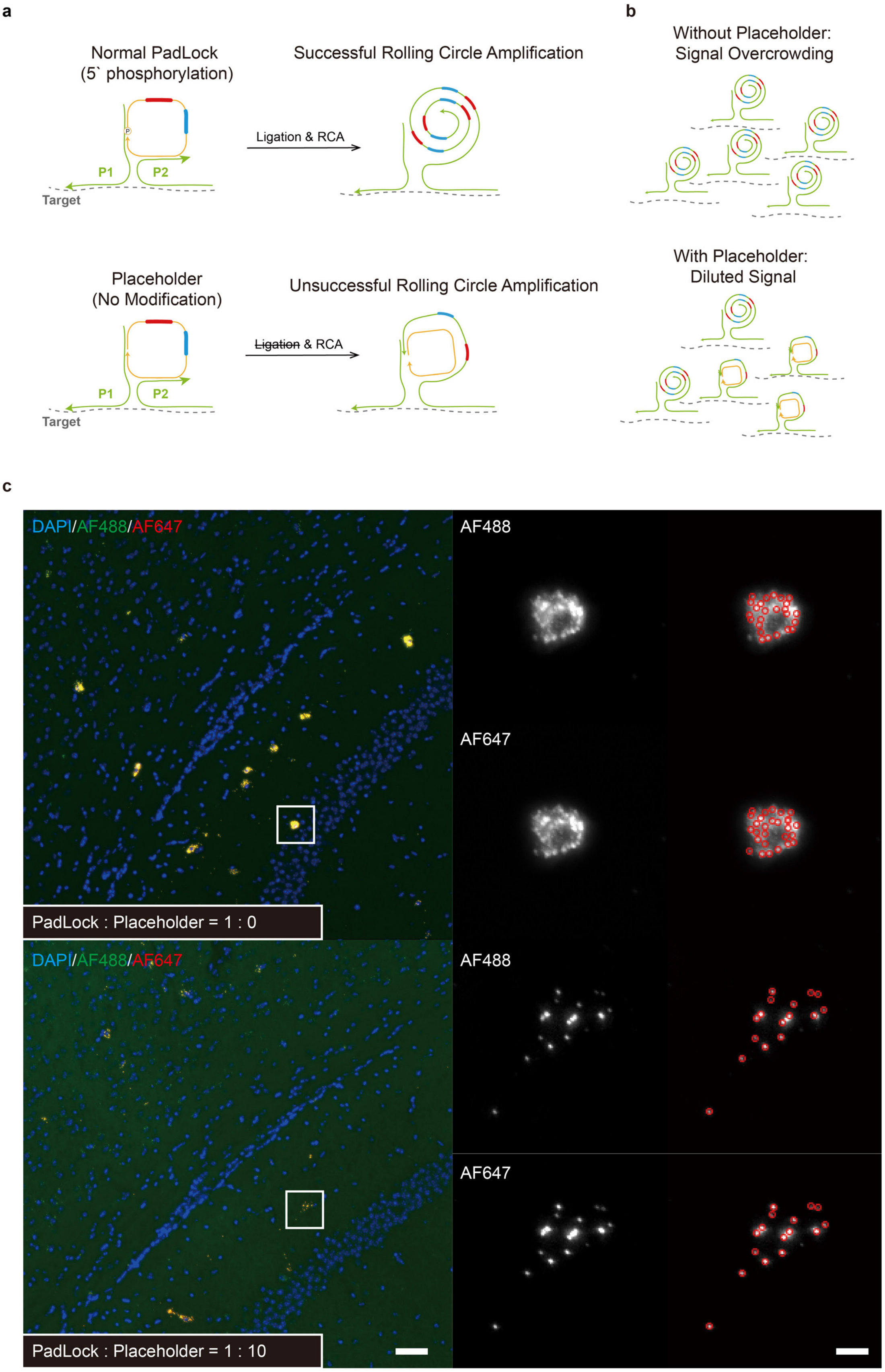
a
Normal PadLock
(5` phosphorylation)
P1
P2
Target
Ligation & RCA
Successful Rolling Circle Amplification
Placeholder
(No Modification)
P1
P2
Target
Ligation & RCA
Unsuccessful Rolling Circle Amplification
b
Without Placeholder:
Signal Overcrowding
With Placeholder:
Diluted Signal
c
DAPI/AF488/AF647
PadLock : Placeholder = 1 : 0
AF488
AF647
DAPI/AF488/AF647
PadLock : Placeholder = 1 : 10
AF488
AF647

**Supplementary Fig. 4: Resolution of Overcrowding. a, b,** Schematic of placeholder mechanism. A mixture of 5'-phosphorylated padlock probes (ligation-competent) and unmodified placeholder probes (ligation-incompetent) was hybridized to target sequences. Only phosphorylated probes undergo ligation and subsequent rolling circle amplification (RCA), while placeholders serve as non-amplifiable competitors. **c,** Demonstration of placeholder effects. The left panels showed *Sst* mRNA detected by OpenFISH, while the right panels displayed corresponding signals identified by RS-FISH. Scale bar left: 200 µm. Scale bar right: 10 µm.

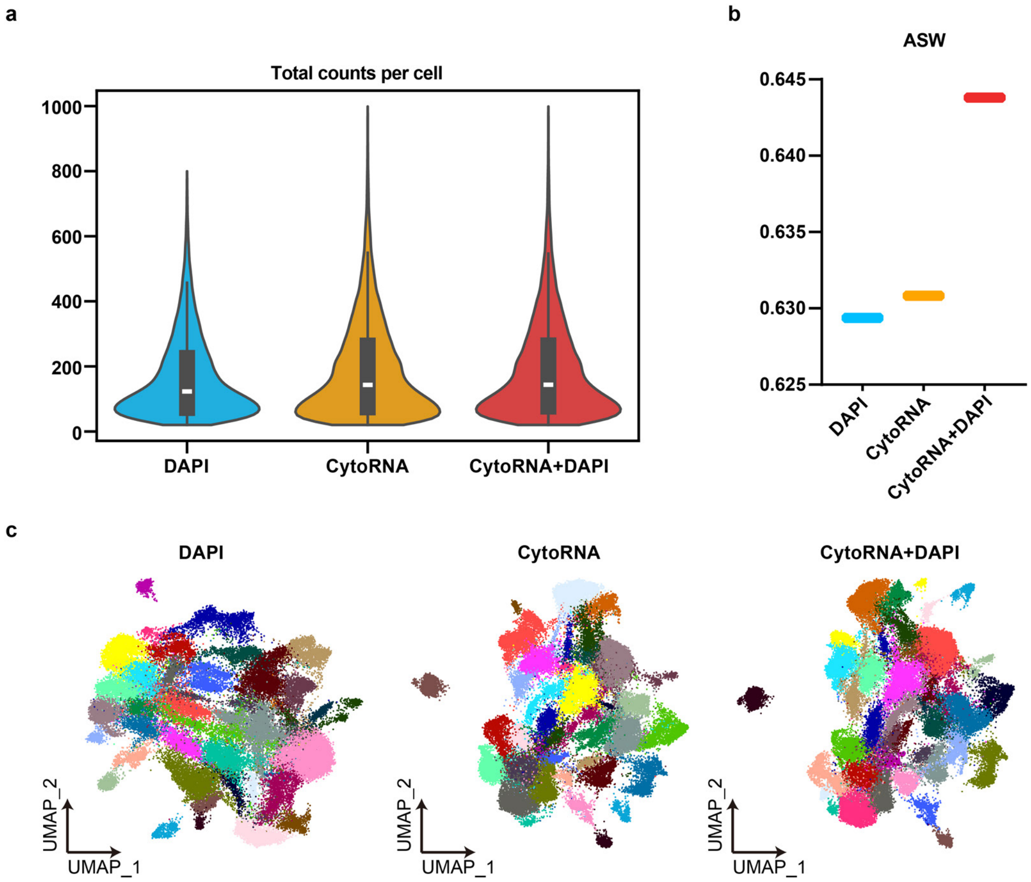

a
Total counts per cell
1000
800
600
400
200
0
DAPI
CytoRNA
CytoRNA+DAPI
b
ASW
0.645
0.640
0.635
0.630
0.625
DAPI
CytoRNA
CytoRNA+DAPI
c
DAPI
CytoRNA
CytoRNA+DAPI
UMAP_2
UMAP_1

**Supplementary Fig. 5: Benchmarking of different segmentation methods. a,** Violin plot displaying total counts per cell for different segmentation modalities used to aggregate data from the OpenFISH mouse brain. **b,** Average silhouette width (ASW) of various segmentation modalities applied to the OpenFISH data, based on the UMAP embeddings derived from unsupervised clustering results **c**. **c,** UMAP visualization of the unsupervised clustering results of OpenFISH mouse brain data under different segmentation modalities.

a

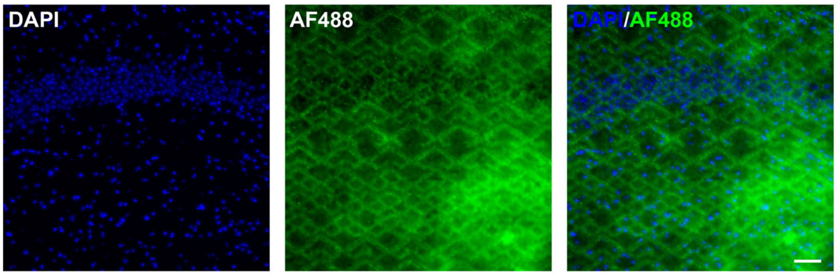
DAPI
AF488
DAPI/AF488

b

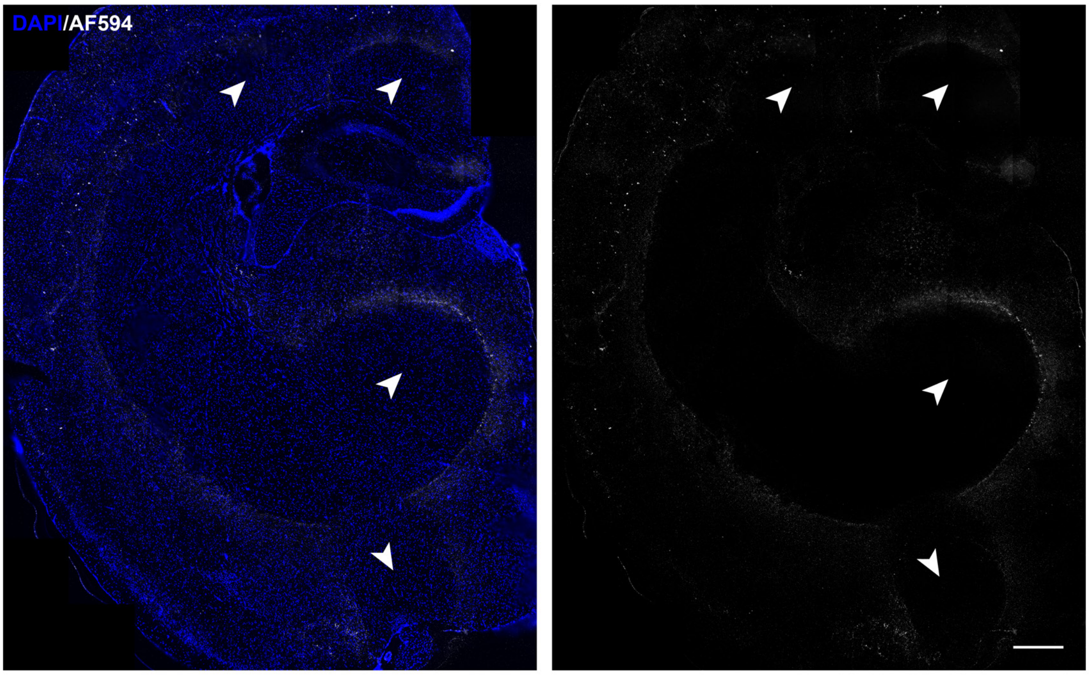
DAPI/AF594

c

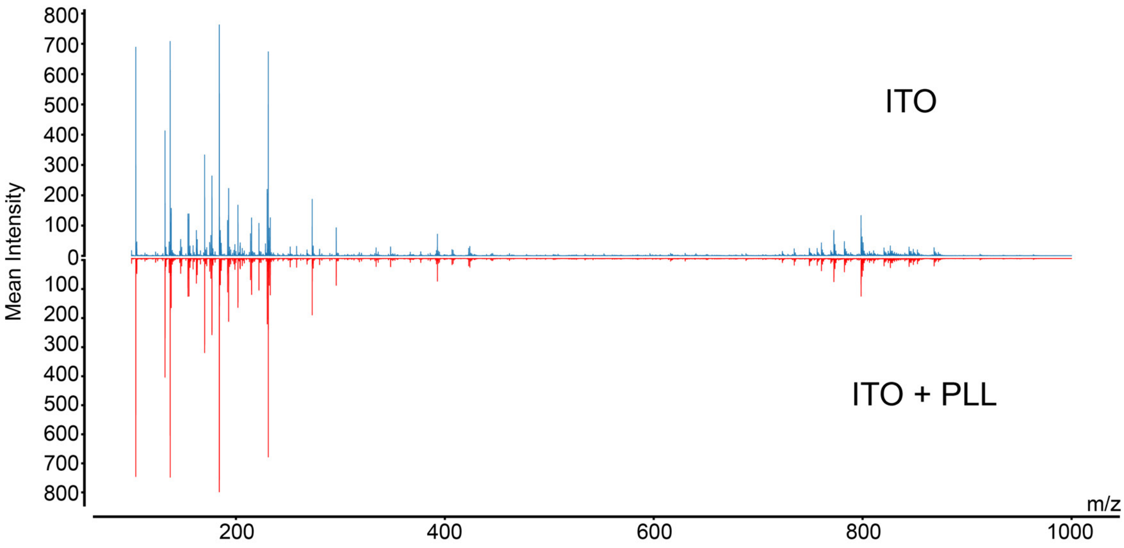
ITO
ITO + PLL
Mean Intensity
m/z
800
700
600
500
400
300
200
100
0
200
400
600
800
1000

**Supplementary Fig. 6: Optimization of OpenFISH protocol for MALDI-MSI integration. a,** Standard OpenFISH protocol performed immediately after MALDI-MSI matrix removal. Scale bar: 100 μm. **b,** After removing the MALDI-MSI matrix, the hydrogel tissue transparency protocol was performed, and then the standard OpenFISH sequencing process was conducted. Scale bar: 500 μm. **c,** The average mass spectra acquired from the tissue sections (top: ITO slide; bottom: PLL-coated ITO slide).

a

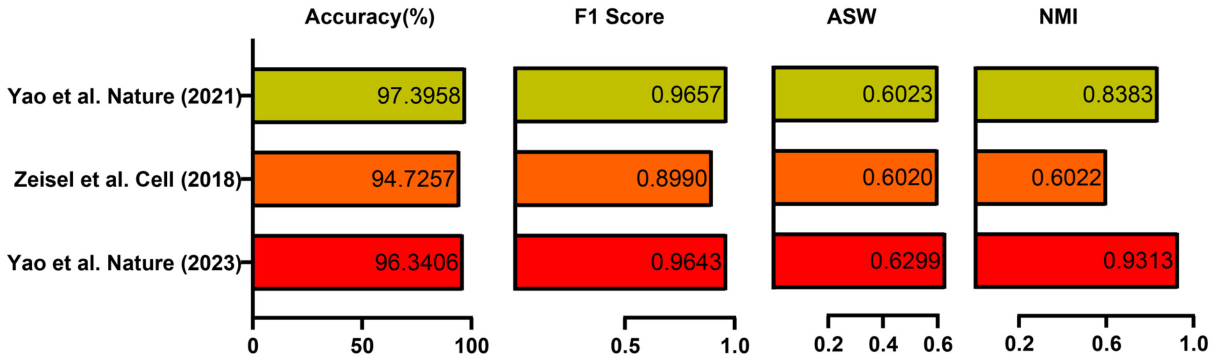
Accuracy(%)
F1 Score
ASW
NMI
Yao et al. Nature (2021)
Zeisel et al. Cell (2018)
Yao et al. Nature (2023)
97.3958
94.7257
96.3406
0.9657
0.8990
0.9643
0.6023
0.6020
0.6299
0.8383
0.6022
0.9313
0
50
100
0.5
1.0
0.2
0.4
0.6
0.2
0.6
1.0

**Supplementary Fig. 7: Benchmarking of panel performance**. **a,** Three scRNA-seq reference datasets were utilized to evaluate the performance of the panel. The ASW was calculated based on UMAP embeddings using prior annotations. The consistency of the unsupervised Leiden clustering results with prior annotations was assessed using normalized mutual information (NMI).

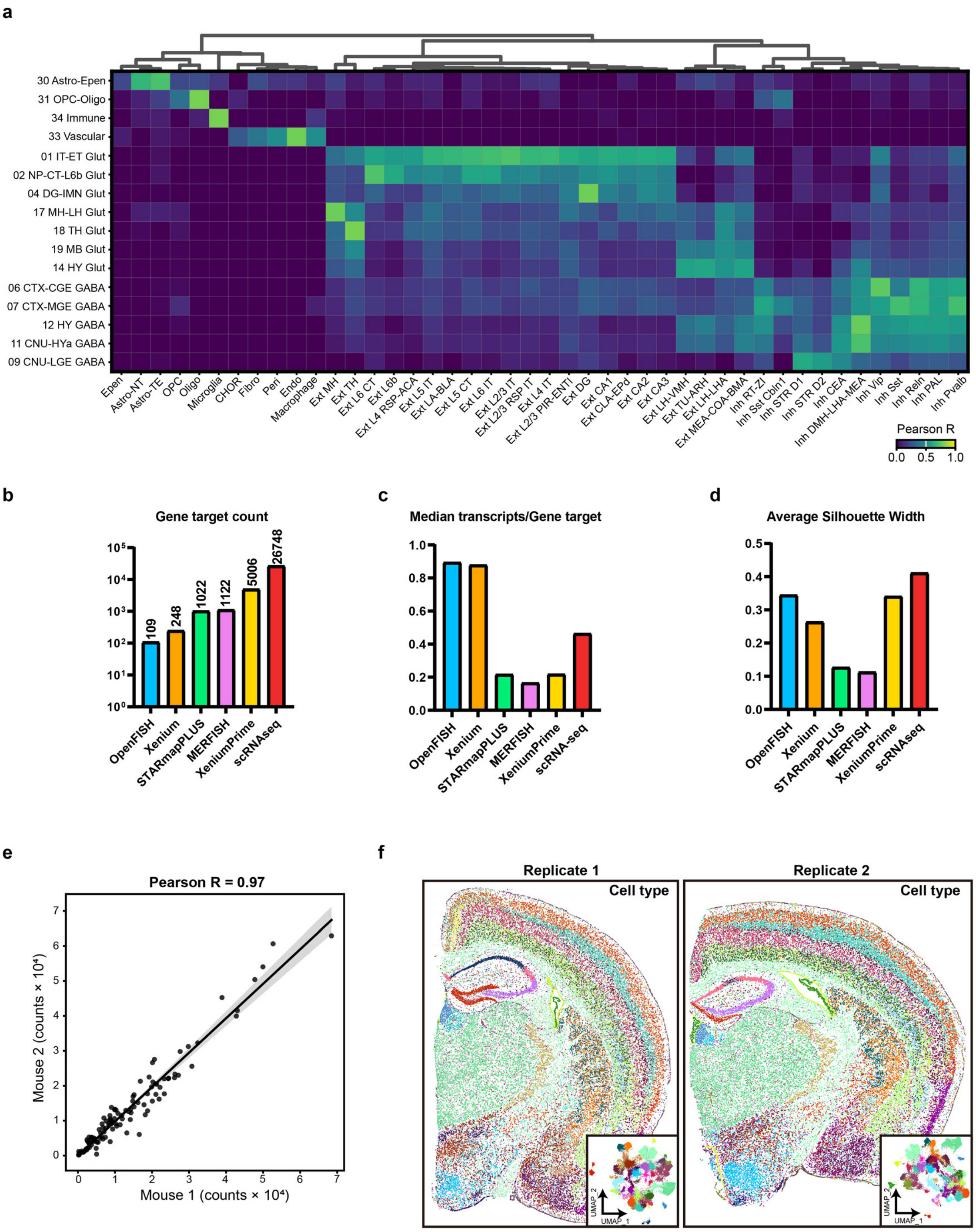
a
30 Astro-Epen
31 OPC-Oligo
34 Immune
33 Vascular
01 IT-ET Glut
02 NP-CT-L6b Glut
04 DG-IMN Glut
17 MH-LH Glut
18 TH Glut
19 MB Glut
14 HY Glut
06 CTX-CGE GABA
07 CTX-MGE GABA
12 HY GABA
11 CNU-HYa GABA
09 CNU-LGE GABA
Pearson R
0.0 0.5 1.0
b
Gene target count
109 248 1022 1122 5006 26748
OpenFISH Xenium STARmapPLUS MERFISH XeniumPrime scRNAseq
c
Median transcripts/Gene target
OpenFISH Xenium STARmapPLUS MERFISH XeniumPrime scRNA-seq
d
Average Silhouette Width
OpenFISH Xenium STARmapPLUS MERFISH XeniumPrime scRNAseq
e
Pearson R = 0.97
Mouse 2 (counts × 10⁴)
Mouse 1 (counts × 10⁴)
f
Replicate 1
Replicate 2
Cell type
UMAP_1
UMAP_2

**Supplementary Fig. 8: Benchmarking of OpenFISH Performance. a,** Cross-reference correspondence of OpenFISH cell types with classes annotated in scRNA-seq datasets across the entire adult mouse brain. **b,** Gene panel size for each method. **c,** The ratio of median transcripts per cell to panel size. **d,** Average silhouette width (ASW) of different methods. **e,** Pearson correlation between two OpenFISH replicates from two replicates. **f,** Cell type annotation of two OpenFISH replicates.

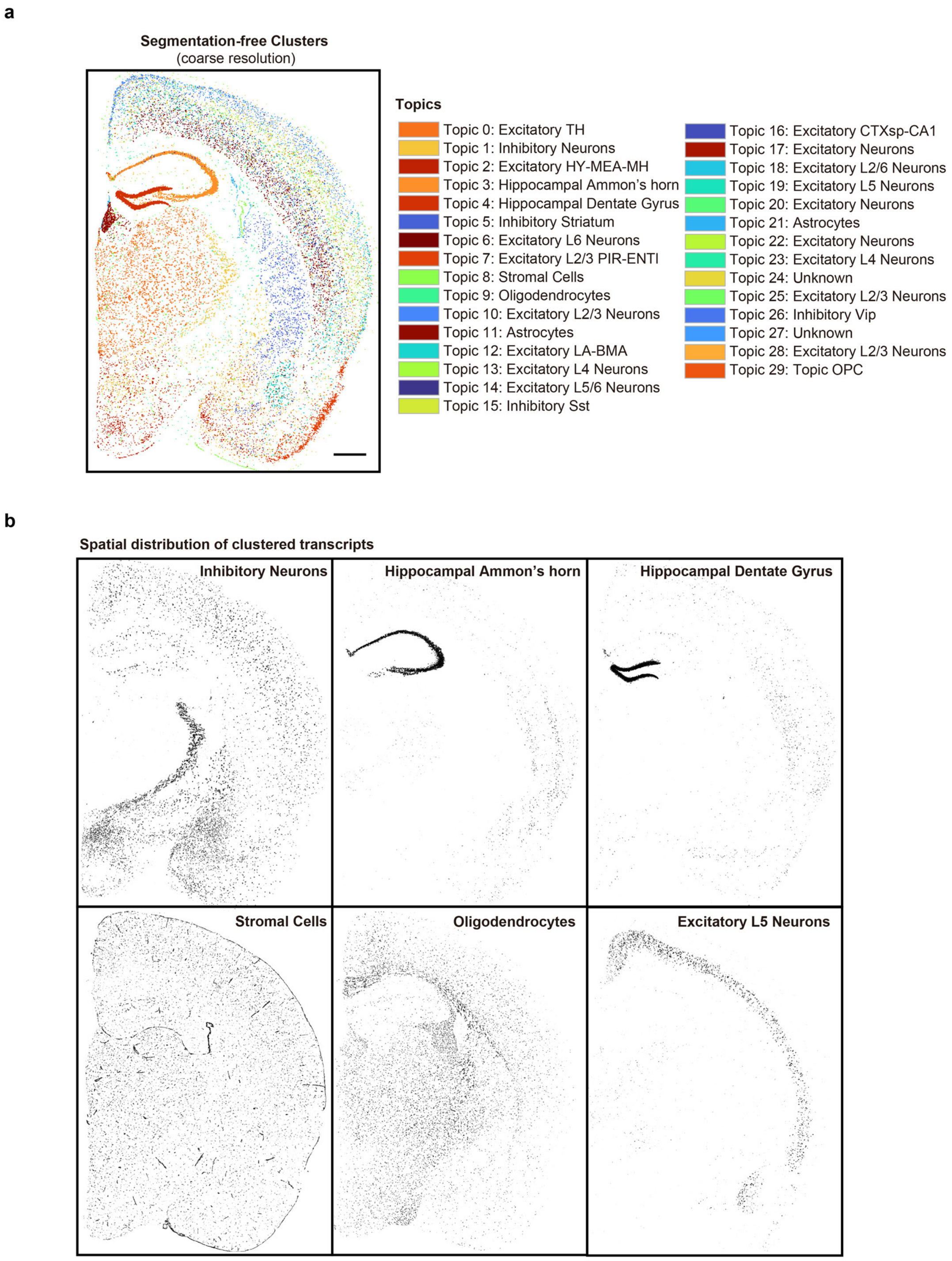
a
Segmentation-free Clusters
(coarse resolution)
Topics
Topic 0: Excitatory TH
Topic 1: Inhibitory Neurons
Topic 2: Excitatory HY-MEA-MH
Topic 3: Hippocampal Ammon's horn
Topic 4: Hippocampal Dentate Gyrus
Topic 5: Inhibitory Striatum
Topic 6: Excitatory L6 Neurons
Topic 7: Excitatory L2/3 PIR-ENTI
Topic 8: Stromal Cells
Topic 9: Oligodendrocytes
Topic 10: Excitatory L2/3 Neurons
Topic 11: Astrocytes
Topic 12: Excitatory LA-BMA
Topic 13: Excitatory L4 Neurons
Topic 14: Excitatory L5/6 Neurons
Topic 15: Inhibitory Sst
Topic 16: Excitatory CTXsp-CA1
Topic 17: Excitatory Neurons
Topic 18: Excitatory L2/6 Neurons
Topic 19: Excitatory L5 Neurons
Topic 20: Excitatory Neurons
Topic 21: Astrocytes
Topic 22: Excitatory Neurons
Topic 23: Excitatory L4 Neurons
Topic 24: Unknown
Topic 25: Excitatory L2/3 Neurons
Topic 26: Inhibitory Vip
Topic 27: Unknown
Topic 28: Excitatory L2/3 Neurons
Topic 29: Topic OPC
b
Spatial distribution of clustered transcripts
Inhibitory Neurons
Hippocampal Ammon's horn
Hippocampal Dentate Gyrus
Stromal Cells
Oligodendrocytes
Excitatory L5 Neurons

**Supplementary Fig. 9: Segmentation-free analysis of OpenFISH. a,** Hexagonal coarse-graining of OpenFISH mouse half brain by FICTURE. Scale bar: 500 μm. **b,** Spatial distribution of selected topics.

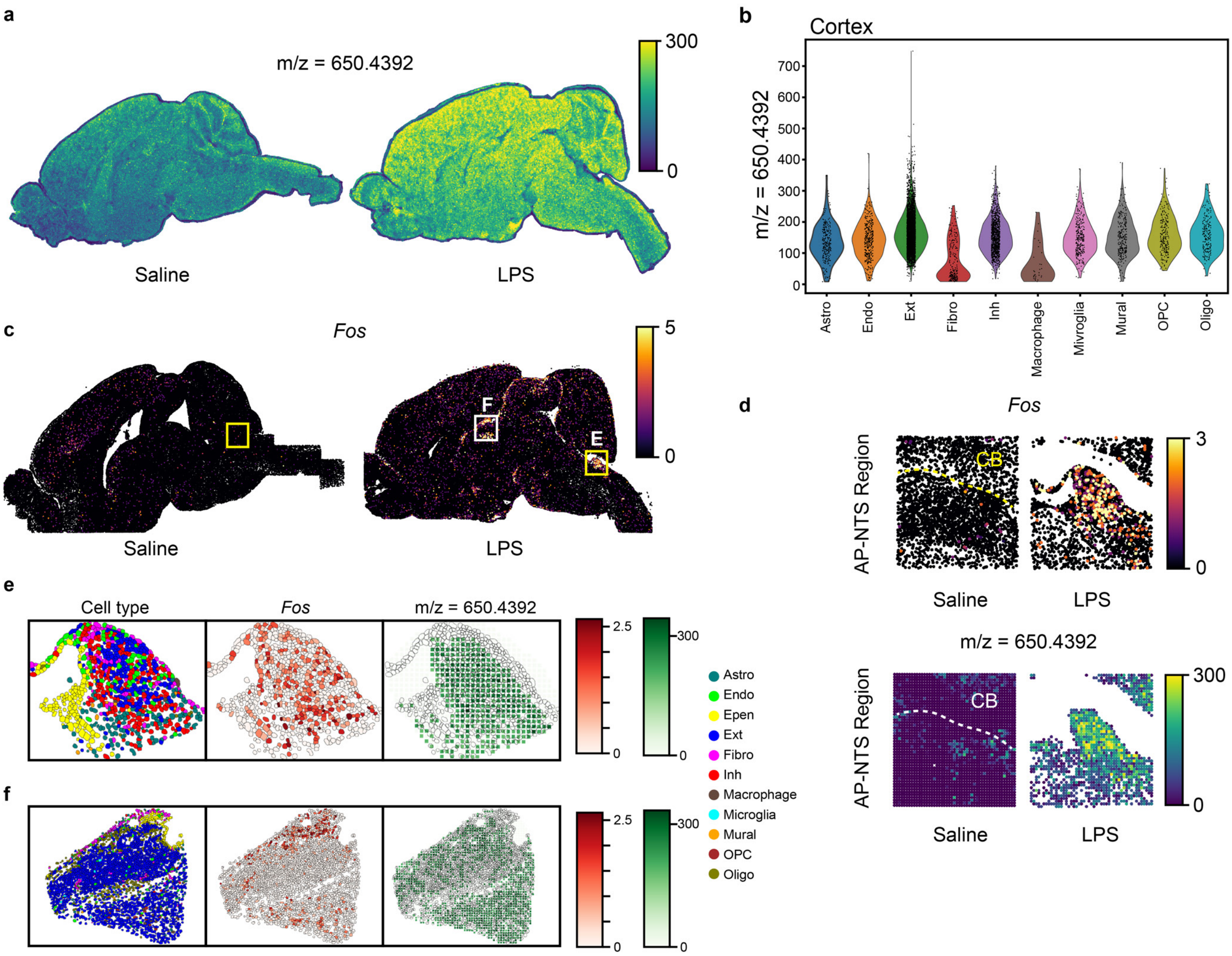
a
m/z = 650.4392
300
0
Saline
LPS
b
Cortex
m/z = 650.4392
0
100
200
300
400
500
600
700
Astro
Endo
Ext
Fibro
Inh
Macrophage
Mivroglia
Mural
OPC
Oligo
c
Fos
5
0
F
E
Saline
LPS
d
Fos
AP-NTS Region
CB
3
0
Saline
LPS
m/z = 650.4392
AP-NTS Region
CB
300
0
Saline
LPS
e
Cell type
Fos
m/z = 650.4392
2.5
0
300
0
Astro
Endo
Epen
Ext
Fibro
Inh
Macrophage
Microglia
Mural
OPC
Oligo
f
2.5
0
300
0

**Supplementary Fig. 10: Molecular Alteration after LPS Treatment. a,** m/z = 650.4392 spatial distribution. **b,** Violin plot of m/z = 650.4392 expression across cell types in cortical region. **c,** *Fos* mRNA spatial distribution. **d,** Magnified region (yellow squared in C) of *Fos* mRNA expression and m/z = 650.4392 intensity. **e,** Cell type distribution, *Fos* mRNA expression and m/z = 650.4392 intensity in selected region. **f,** Cell type distribution, *Fos* mRNA expression and m/z = 650.4392 intensity in selected region.

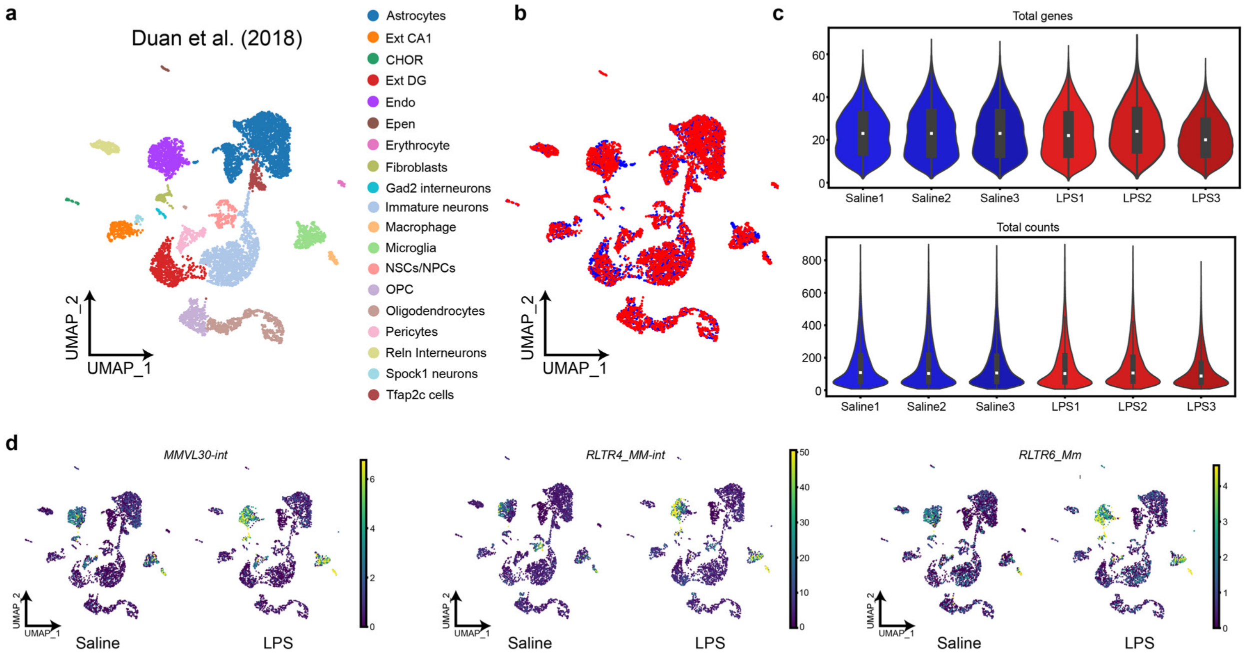
a
Duan et al. (2018)
Astrocytes
Ext CA1
CHOR
Ext DG
Endo
Epen
Erythrocyte
Fibroblasts
Gad2 interneurons
Immature neurons
Macrophage
Microglia
NSCs/NPCs
OPC
Oligodendrocytes
Pericytes
Reln Interneurons
Spock1 neurons
Tfap2c cells
UMAP_1
UMAP_2
b
c
Total genes
Total counts
Saline1
Saline2
Saline3
LPS1
LPS2
LPS3
d
MMVL30-int
RLTR4_MM-int
RLTR6_Mm
Saline
LPS

**Supplementary Fig. 11: Cell-type-specific transcriptional responses to inflammation. a,** Cell type annotation of single-cell RNA sequencing data from Duan et al. (2018). NSCs: Neural Stem cells; NPCs: Neural Progenitor cells. **b,** Evaluation of dataset integration quality between LPS-treated and saline-treated conditions. **c,** Quality control metrics for OpenFISH experiments. **d,** Expression profiles of transposable element (TE) subfamilies in the scRNA-seq reference dataset, matched to OpenFISH targets.

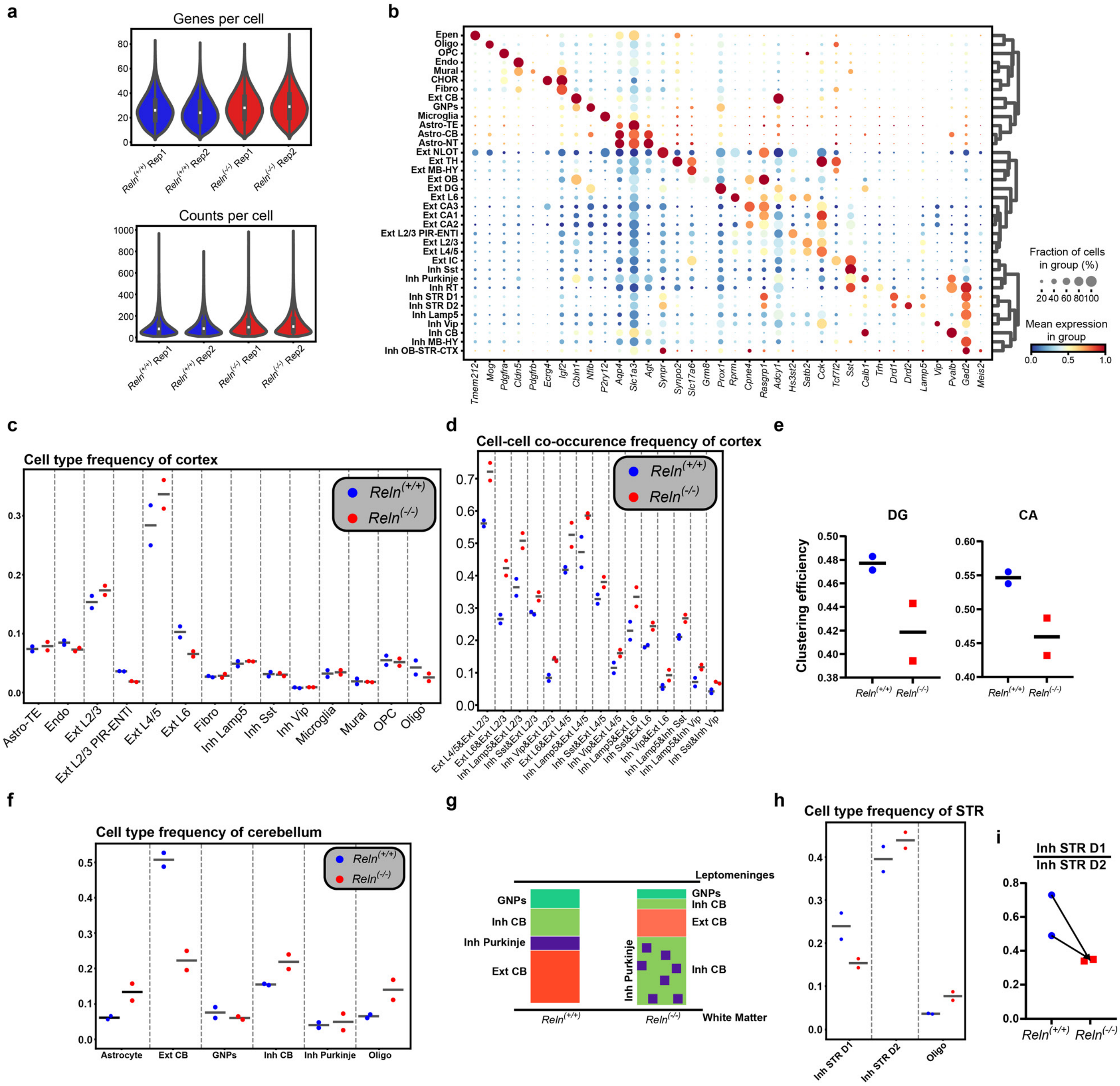
a
Genes per cell
Counts per cell
Reln(+/+) Rep1
Reln(+/+) Rep2
Reln(-/-) Rep1
Reln(-/-) Rep2
b
Epen
Oligo
OPC
Endo
Mural
CHOR
Fibro
Ext CB
GNPs
Microglia
Astro-TE
Astro-CB
Astro-NT
Ext NLOT
Ext TH
Ext MB-HY
Ext OB
Ext DG
Ext L6
Ext CA3
Ext CA1
Ext CA2
Ext L2/3 PIR-ENTl
Ext L2/3
Ext L4/5
Ext IC
Inh Sst
Inh Purkinje
Inh RT
Inh STR D1
Inh STR D2
Inh Lamp5
Inh Vip
Inh CB
Inh MB-HY
Inh OB-STR-CTX
Tmem212 Mog Pdgfra Cldn5 Pdgfrb Ecrg4 Igf2 Cbln1 Nfib P2ry12 Aqp4 Slc1a3 Agt Synpr Synpo2 Slc17a6 Grm8 Prox1 Rprm Cpne4 Rasgrp1 Adcy1 Hs3st2 Satb2 Cck Tcf7l2 Sst Calb1 Trh Drd1 Drd2 Lamp5 Vip Pvalb Gad2 Meis2
Fraction of cells in group (%)
20 40 60 80 100
Mean expression in group
0.0 0.5 1.0
c
Cell type frequency of cortex
Reln(+/+)
Reln(-/-)
Astro-TE
Endo
Ext L2/3
Ext L2/3 PIR-ENTl
Ext L4/5
Ext L6
Fibro
Inh Lamp5
Inh Sst
Inh Vip
Microglia
Mural
OPC
Oligo
d
Cell-cell co-occurence frequency of cortex
Reln(+/+)
Reln(-/-)
Ext L4/5&Ext L2/3
Ext L6&Ext L2/3
Inh Lamp5&Ext L2/3
Inh Sst&Ext L2/3
Inh Vip&Ext L2/3
Ext L6&Ext L4/5
Inh Lamp5&Ext L4/5
Inh Sst&Ext L4/5
Inh Vip&Ext L4/5
Inh Lamp5&Ext L6
Inh Sst&Ext L6
Inh Vip&Ext L6
Inh Lamp5&Inh Sst
Inh Lamp5&Inh Vip
Inh Sst&Inh Vip
e
DG
CA
Clustering efficiency
Reln(+/+)
Reln(-/-)
f
Cell type frequency of cerebellum
Reln(+/+)
Reln(-/-)
Astrocyte
Ext CB
GNPs
Inh CB
Inh Purkinje
Oligo
g
Leptomeninges
GNPs
Inh CB
Inh Purkinje
Ext CB
Inh Purkinje
White Matter
Reln(+/+)
Reln(-/-)
h
Cell type frequency of STR
Inh STR D1
Inh STR D2
Oligo
i
Inh STR D1 / Inh STR D2
Reln(+/+)
Reln(-/-)

**Supplementary Fig. 12: Structural abnormalities in *Reln*$^{(-/-)}$ neurodevelopmental model. a,** Quality control of *Reln*$^{(+/+)}$ and *Reln*$^{(-/-)}$ samples **b,** Dot plot displaying marker gene expression for each cell type. **c,** Relative frequency of cell types in the cortical region. **d,** Co-occurrence frequency of cell-cell interactions in the cortical region. **e,** Clustering efficiency of excitatory neurons (Ext DG and Ext CA subtypes) in the hippocampal region. **f,** Cell type composition in the cerebellar region. **g,** Schematic illustration of structural abnormalities in the cerebellar region. **h,** Distribution of cell types in the striatal region. **i,** Ratio of D1- to D2-expressing inhibitory neurons (Inh STR D1/D2) in the striatal region.
